# Nausea-induced suppression of feeding is mediated by central amygdala Dlk1 expressing neurons

**DOI:** 10.1101/2023.06.29.547043

**Authors:** Wenyu Ding, Helena Weltzien, Christian Peters, Rüdiger Klein

**Author notes:** Correspondence (R.K.).

## Abstract

The motivation to eat is suppressed by satiety and by aversive stimuli such as nausea. Compared to the neural regulation of homeostatic feeding, the mechanism of appetite suppression by nausea is not well understood. Previous work characterized PKCδ neurons in the lateral subdivision (CeL) of the central amygdala (CeA) to suppress feeding in response to satiety signals and nausea. Here, we characterized a previously unknown neuronal population enriched in the medial subdivision (CeM) of the CeA and marked by expression of Dlk1. Distinct from CeA^PKCδ^ neurons, CeA^Dlk1^ neurons are activated by nausea, but not by satiety, form long-range projections to many brain regions and exert their anorexigenic activity by inhibition of neurons of the parabrachial nucleus. CeA^Dlk1^ neurons are under inhibitory control of appetitive CeA neurons, but also receive long-range monosynaptic inputs from multiple brain regions. Our results illustrate a novel CeA circuit that regulates nausea-induced feeding suppression.

**Highlights:** CeA^Dlk1^ neurons are a previously unknown CeA cell population, enriched in the CeM

CeA^Dlk1^ neurons are activated by nausea and bitter food, but not satiety

CeA^Dlk1^ neurons suppress feeding under conditions of nausea

CeA^Dlk1^ neuronal projections to the PBN mediate feeding suppression

## Introduction

Organisms suppress their feeding behavior to adapt to potentially risky internal and external factors in order to survive. For example, animals can maintain energy homeostasis by secreting anorexigenic hormones like amylin and cholecystokinin (CCK), which suppress appetite ^1, 2^. Additionally, exogenous compounds such as lithium chloride (LiCl), a salt that creates gastric discomfort, and lipopolysaccharide (LPS), a bacterial cell wall component that induces inflammation, have appetite-suppressing effects ^3^. Feeding behavior can also be suppressed by emotional states such as stress and anxiety. The central amygdala (CeA) is thought to be involved in regulating emotional feeding behavior, and evidence suggests that the CeA is an important nucleus in regulating feeding behavior during aversive conditioning ^4^.

The CeA is a striatal-like structure organized into capsular, lateral, and medial subdivisions (CeC, CeL, and CeM) and composed of multiple discrete GABAergic neurons. The CeL receives multimodal inputs, processes the information, and produces scaled outputs in part via the CeM, which has long-range projections to many brain regions ^5^. As previously shown, a population in the CeL labelled by protein kinase C-delta (Pkcδ^+^) mediates the effects of several anorexigenic signals, such as satiety, nausea and bitter taste. Silencing of CeA^Pkcδ^ neurons can largely block the eating inhibition elicited by these anorexigenic signals, and activation of CeA^Pkcδ^ neurons is sufficient to suppress food intake ^3^. CeA^Pkcδ^ neurons in the CeC region, labelled by Calcrl, are involved in suppressing food intake by receiving projections from calcitonin gene related peptide (CGRP) neurons in the lateral parabrachial nucleus (lPBN), which transmit anorexigenic and danger signals to the CeA ^6, 7^. CeA^Pkcδ^ neurons form efferent projections to the bed nucleus of the stria terminalis (BNST), a brain area known to be involved in negative emotions. However, studies using optogenetics to activate the terminals in the BNST that receive input from CeA^Pkcδ^ neurons did not result in a suppression of food intake ^3^. This suggests that this pathway is unlikely to play a major role in regulating feeding behavior. Another target of CeA^Pkcδ^ neuron projections is the substantia innominata (SI), which is the primary source of cortical cholinergic innervation ^8^. By projecting to the SI, these neurons are thought to mediate negative reinforcement learning. Nonetheless, this pathway is also unlikely to be the primary output for suppressing feeding behavior ^9^. It is likely that CeA^Pkcδ^ neurons elicit their anorexigenic effects by inhibiting local appetitive neurons ^3^, which brings up the question of how information from the CeA is transmitted to other brain regions. The suppression of feeding should involve many brain regions, such as the arcuate nucleus (Arc) and lateral hypothalamus (LH) that regulate appetite ^10, 11^, as well as the PBN and paraventricular nucleus of the thalamus (PVT) that process aversive stimuli ^12, 13^, Surprisingly, these brain regions are not among the outputs of CeA^Pkcδ^ neurons.

Recent studies using single-cell RNA sequencing (scRNA-seq) and spatial profiling have improved our understanding of the cell type organization of the CeA ^14–16^. Previously unknown molecular cell types were characterized, enriched in the CeM, such as the Delta like non canonical Notch ligand 1 (Dlk1) and vitamin D receptor (Vdr) populations ^14^, as well as the Nr2f2 and Isl1 clusters ^15^. Many long-range CeA projections are associated with these previously unresolved molecular cell types raising the possibility that one or more of these CeM clusters may have anorexigenic functions.

The PBN is another brain region that is thought to mediate the suppression of appetite induced by anorectic hormones like amylin and CCK, as well as by LiCl and LPS. It serves as a major hub for sensory information relevant to food and water intake, nociceptive response, malaise, and aversive emotional behaviors, receiving inputs from both interoceptive and exteroceptive sources ^6, 7, 12, 17–20^. The PBN is comprised of over 10 transcript-defined glutamatergic neuron types expressing an abundance of different neuropeptides and neuropeptide receptors ^19^. Among them, CGRP neurons in the lPBN respond to negative visceral stimuli, such as LiCl and project their axons to the CeA ^18^. Conversely, neurons in the CeA send GABAergic inputs to the PBN, typically to the lPBN, to mediate appetitive or reward behavior. For example, serotonin receptor 2a (Htr2a)-expressing neurons in the CeA promote feeding and positive reinforcement by inhibiting neurons in the PBN ^21^. These neurons overlap with prepronociceptin-positive (Pnoc) neurons ^22^, which also project to the PBN, producing reward behavior ^23^. Additionally, CeA neurons expressing neurotensin (NTS) project to the PBN, increasing the consumption of rewarding and/or palatable fluids ^24^.

The current study aimed at investigating whether a specific population of cells in the CeA could transmit information to distant brain regions and contribute to the suppression of feeding behaviors. We focused our attention on the newly identified CeA^Dlk1^ cluster ^14^, whose transcriptome is most similar to CeA^Pkcδ^ neurons, and which is predominantly located in the CeM region. Distinct from CeA^Pkcδ^ neurons, CeA^Dlk1^ neurons are activated by nausea-inducing agents, but not by satiety, form long-range inhibitory projections to many brain regions and suppress food intake at least in part by inhibiting neurons of the PBN. The activity of CeA^Dlk1^ neurons is controlled by local presynaptic input from appetitive CeA neurons. In addition, CeA^Dlk1^ neurons receive distinct monosynaptic inputs from brain regions linked to sensory experience, odors, and aversive stimuli. Our investigation has revealed critical neural players and circuit mechanisms that regulate nausea-induced feeding suppression.

## Results

### CeA^Dlk1^ neurons are a previously unknown cell population in the CeA

From CeA snRNA-seq datasets, we and others identified a previously unknown cell cluster marked by expression of Dlk1 ^14, 22^. To compare the transcriptome of this new cluster with other well-known CeA populations, we subclustered the snRNA-seq data from two data sets from 8-week old naïve mice ^22^, which included two Pkcδ-positive clusters (one residing in the CeL, CeL^Pkcδ^, and one residing in the CeC, CeC^Cdh^^9.Calcrl^), two Sst-positive clusters (one residing in the CeL, CeL^Sst^, and one residing in the CeM, CeM^Tac^^1.Sst^), and the CeA^Dlk1^ cluster. The t-distributed stochastic neighbor embedding (TSNE) visualization (Figure 1A) and differently expressed genes (DEGs) (Figure 1B) suggested that CeA^Dlk1^ neurons comprise a distinct cell type expressing cell-type-specific genes. We constructed a taxonomy tree (Figure 1C) based on the correlation of the expression of highly variable genes (VGs) across cell types. The pairwise correlation heatmaps of each cluster revealed that the two Pkcδ-positive clusters displayed high correlation. The CeA^Dlk1^ cluster showed week correlation with the CeL^Sst^, moderate correlation with the CeM^Tac^^1.Sst^, and high correlation with the CeL^Pkcδ^ cluster. These results suggest that the CeA^Dlk1^ cluster is a distinct cell population with more similarities to CeA^Pkcδ^ than the two CeA^Sst^ clusters.

**Figure 1.**
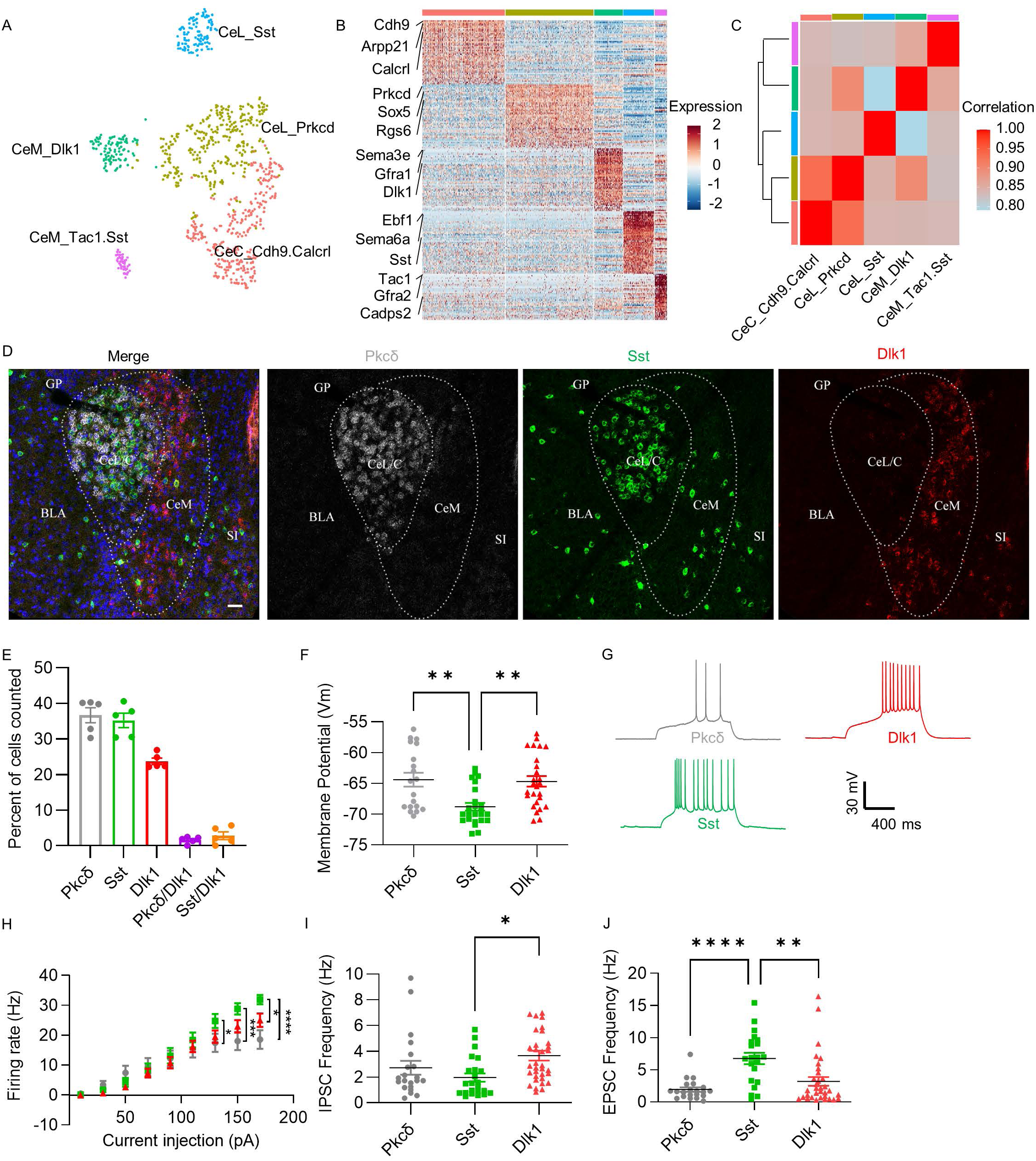
CeA^Dlk1^ neurons, a newly identified cell population in the CeA. **A** t-distributed stochastic neighbor embedding (t-SNE) representation of neurons in the CeA colored by cluster. **B** A heatmap showing the expression of the top 50 differentially expressed markers of each cluster across all cells. Color code as in panel A. **C** Unsupervised hierarchical clustering of pairwise Pearson correlation coefficients of transcripts in RNA-Seq analysis. **D** Representative image of Dlk1 (red), Sst (green) and Pkcδ (grey) mRNA expression in the CeA. Scale bar, 30 μm. **E** Quantification of Pkcδ, Dlk1, Sst, and double-positive cell counts across all CeA images (4 sections per mouse, 5 mice; error bars represent standard error of the mean (SEM)). **F-J** Physiological properties of the three cell populations characterized in acute slices. F Quantifications of membrane potentials (Vm), n=19 neurons for Pkcδ, n=24 neurons for Sst, n=26 neurons for Dlk1. One-way ANOVA, **p<0.01. **G** Representative whole-cell current-clamp recordings. **H** Firing rates (Hz) after injecting different current steps. n=18 neurons for Pkcδ, n=26 neurons for Sst, n=26 neurons for Dlk1. One-way ANOVA, *p<0.05, ***p<0.001, ****p<0.0001. **I** Quantification of sIPSC frequency, n=21 neurons for Pkcδ, n=23 neurons for Sst, n=34 neurons for Dlk1. One-way ANOVA, *p<0.05. **J** Quantification of sEPSC frequency, n=21 neurons for Pkcδ, n=23 neurons for Sst, n=34 neurons for Dlk1. One-way ANOVA, **p<0.01, ****p<0.0001.

To gain a better understanding of the spatial relationship between CeA^Dlk1^, CeA^Sst^ and CeA^Pkcδ^ neurons, we performed fluorescence in situ hybridization (FISH) with Dlk1, Sst, and Pkcδ probes (Figures 1D and S1A). We observed a large population of Dlk1-positive cells in the CeM and only few cells in the CeL/C. Dlk1-positive cells overlapped weakly with Pkcδ (1.5%), and Sst-positive cells (2.8%) (Figure 1E). We performed similar dual FISH with Dlk1 and Crh, Drd2, Pnoc, Tac2, and Nts (Figure S1B-K). Across all of these markers, Dlk1-positive cells had weak overlapping expression with Crh (6.3%), Pnoc (5.7%), Tac2 (3.8%), and Nts (6.0%), and moderate overlap with Drd2 (21.4%). To characterize the overlap with Htr2a-positive cells, we bred Htr2a-Cre mice with a LacZ reporter line and then performed ISH of Dlk1 combined with β-Gal immunostaining in Htr2a-Cre::LacZ mice. Under these conditions, we observed weak overlap (0.7%) (Figure S1L,M). Together, these results support the notion that CeA^Dlk1^ neurons are a previously unknown CeA cell population, enriched in the CeM, with little spatial overlap with other CeA cell types.

To investigate the role of CeA^Dlk1^ neurons in appetitive behavior, we used Dlk1-CreER mice ^25^. To verify the fidelity of Cre expression in this mouse line, we performed dual fluorescence in situ hybridization for LacZ and Dlk1 in the CeA of Dlk1-CreER::LacZ mice, by breeding Dlk1-CreER mice with a LacZ reporter line (Figure S2A). We observed detectable Dlk1 expression in > 77 % of the LacZ-expressing cells (Figure S2B). Thus, we determined that the Dlk1-CreER mouse is a robust, high-fidelity strain by which to gain genetic access to Dlk1-expressing cells in the CeA.

To describe the physiological properties of Dlk1-expressing cells in the CeA, we used ex vivo slice patch-clamp electrophysiology in Dlk1-CreER::tdTomato mice. We also patched CeA^Sst^ and CeA^Pkcδ^ neurons to compare the physiological properties of CeA^Dlk1^ neurons to mainly appetitive (Sst) and aversive (Pkcδ) cell populations. The results suggest that the membrane potential of CeA^Dlk1^ neurons was quite different from CeA^Sst^, but similar to CeA^Pkcδ^ neurons (Figure 1F). In terms of basic firing rates, CeA^Sst^ neurons exhibited a higher frequency than CeA^Pkcδ^ neurons, as previously shown ^26^. We found that the CeA^Dlk1^ neuron firing rate was somewhere in between these two populations (Figure 1G, H).

The frequencies of spontaneous inhibitory postsynaptic currents (sIPSC) were higher in CeA^Dlk1^ than CeA^Sst^ neurons, suggesting that CeA^Dlk1^ neurons received more inhibitory inputs compared to CeA^Sst^ neurons (Figure 1I). This result was consistent with the anatomic location of CeA^Dlk1^ neurons being mainly in CeM, where they might be receiving dense inhibition from CeL ^27^. In contrast, spontaneous excitatory postsynaptic current (sEPSC) frequencies were higher in CeA^Sst^ compared to CeA^Dlk1^ and CeA^Pkcδ^ neurons, suggesting that CeA^Sst^ neurons receive more excitatory inputs than the other populations (Figure 1J). These results suggest that CeA^Dlk1^ neurons have unique intrinsic physiological properties, that are more similar to aversive (CeA^Pkcδ^) than appetitive (CeA^Sst^) neurons.

### CeA^Dlk1^ neurons are activated by nausea and bitter food

Since CeA^Dlk1^ neurons have a similar transcriptome and physiological properties as CeA^Pkcδ^ neurons, we hypothesized that CeA^Dlk1^ neurons may also be activated by anorexigenic signals. We monitored c-Fos expression after intraperitoneal injection of cholecystokinin (CCK), a peptide which mimics satiety ^28^, and lithium chloride (LiCl), which induces nausea and visceral malaise ^29^. Intraperitoneal injection of saline was used as control. Immunostaining for c-Fos revealed that CCK induced clear c-Fos expression in the CeA, and that this effect was mostly due to c-Fos expression in Dlk1-negative neurons. In contrast, LiCl injection induced strong c-Fos expression in the CeA, including a sizeable fraction of CeA^Dlk1^ neurons (Figure 2A-D).

**Figure 2.**
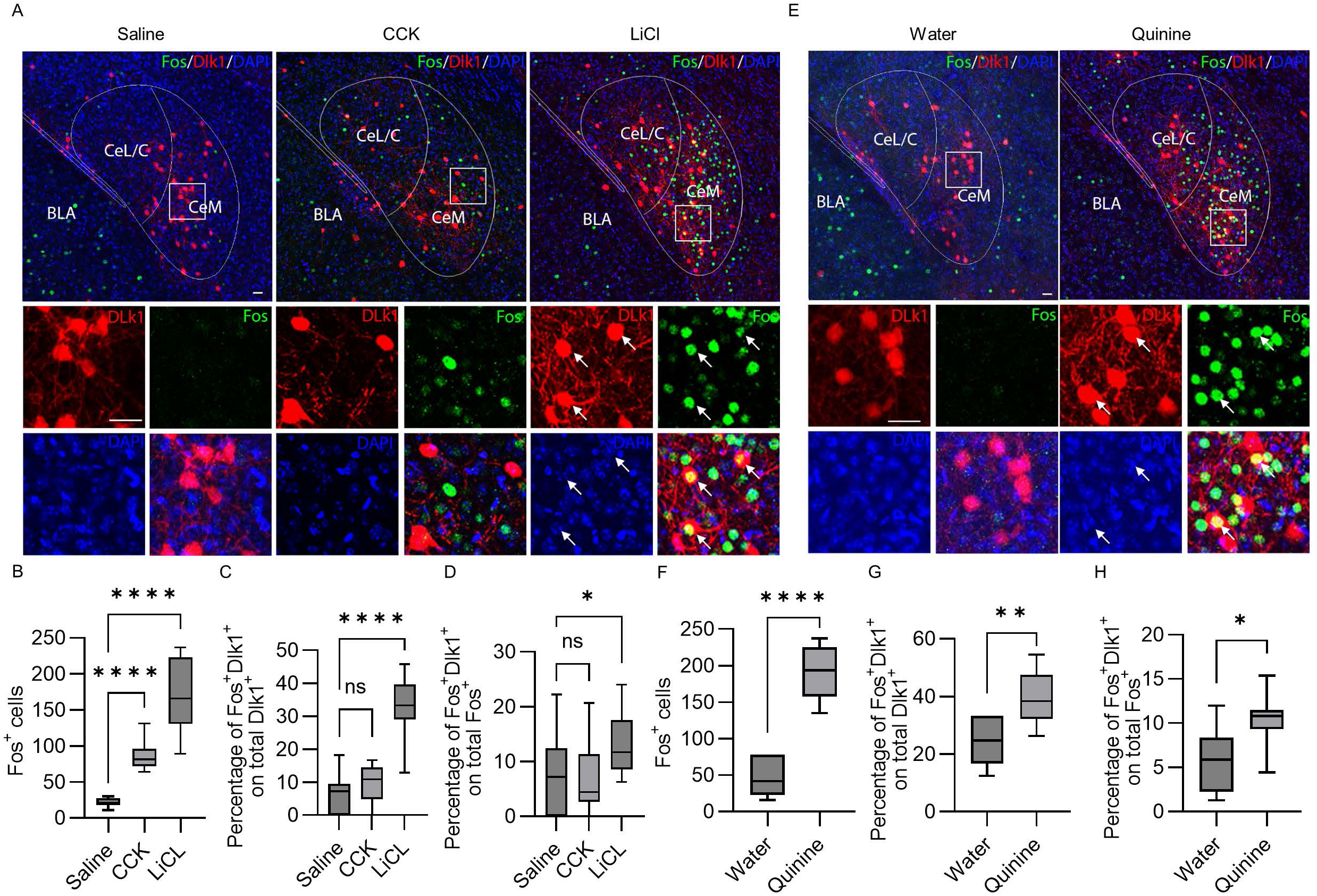
CeA^Dlk1^ neurons are activated by LiCl and quinine. **A** Representative images of c-Fos immunostainings (green) in CeA of Dlk1-CreER::Ai9 tdTomato mice (red) after intraperitoneal injection of saline, CCK or LiCl. White box indicates the location of the high-magnification panel on the bottom. Arrows indicate double-positive cells. Scale bars, 30 μm. **B** Percentages of c-Fos^+^ cells in CeA after the indicated treatments. Data in panels B-D were analyzed by One-way ANOVA (n=3 mice in each condition). For saline versus CCK, P< 0.0001. For saline versus LiCl, P < 0.0001. **C** Percentages of c-Fos^+^/Dlk1^+^ cells among Dlk1^+^ cells in CeA after the indicated treatments. For saline versus CCK, P = 0.3611. For saline versus LiCl, P < 0.0001. **D** Percentages of c-Fos^+^/Dlk1^+^ cells among c-Fos^+^ cells in CeA after the indicated treatments. For saline versus CCK, P = 0.9468. For saline versus LiCl, P = 0.0285. **E** Representative images of c-Fos immunostainings (green) in CeA of Dlk1-CreER::Ai9 tdTomato mice (red) after oral infusion of water or quinine solution. White box indicates the location of the high-magnification panel on the bottom. Arrows indicate double-positive cells. **F** Percentages of c-Fos^+^ cells in CeA after the indicated treatments. Data in panels F-H were analyzed by Two-tailed unpaired t test, P < 0.0001. **G** Percentages of c-Fos^+^/Dlk1^+^ cells among Dlk1^+^ cells in CeA after the indicated treatments. P = 0.0048. H Percentages of c-Fos^+^/Dlk1^+^ cells among c-Fos^+^ cells in CeA after the indicated treatments. P = 0.0149.

To further investigate whether CeA^Dlk1^ neurons became activated during a normal feeding cycle, we compared c-Fos expression between fasted mice that had been food deprived for 20h, and ad libitum fed control mice. We found that food deprivation strongly induced c-Fos expression in CeA compared to satiety (Figure S3A, B) (see also ^30^). In line with the CCK results, the numbers of c-Fos^+^ cells within the CeA^Dlk1^ cluster did not change between fasted and satiated mice (Figure S3C, D). Next, we examined activation of CeA^Dlk1^ neurons by the bitter tastant quinine, which also reduces food intake. We observed an increase in c-Fos expression in the CeA after oral infusion of quinine, compared to water. This increase was, in part, due to an increase in cFos^+^ CeA^Dlk1^ neurons (Figure 2E-H). These results suggest that a sizeable fraction of CeA^Dlk1^ neurons was activated by nausea and unpalatable liquids, but not by interoceptive signals associated with satiety.

### Activation of CeA^Dlk1^ neurons inhibits feeding

Next, we used optogenetic stimulation experiments to assess the functions of CeA^Dlk1^ neurons. Since CeA^Dlk1^ neurons were activated by anorexigenic agents, we hypothesized that photoactivation of CeA^Dlk1^ neurons may be sufficient to inhibit food intake. We injected a Cre-dependent AAV expressing ChR2 bilaterally into the CeA of Dlk1-CreER mice, induced Cre expression by Tamoxifen and first confirmed using whole-cell patch-clamp recordings in CeA brain slices that 473-nm light pulses triggered robust spiking in CeA^Dlk1^ neurons with up to 20Hz stimulation (Figure 3A, B). For feeding assays, we implanted optical fibers bilaterally over the CeA (Figure 3C and S4A) and measured food intake for 30 min in fasted mice, first with photoactivation of CeA^Dlk1^ neurons, followed by another 30 min in the absence of light (Figure 3D). We found that photoactivation of CeA^Dlk1^ neurons expressing ChR2 strongly inhibited food intake, compared to photoactivated mice expressing a Cre-dependent EGFP virus (Figure 3E). In the subsequent ‘light off’ phase, ChR2-expressing mice resumed feeding, whereas control mice significantly decreased food intake (Figure 3E). A similar inhibition of food intake was observed in photoactivated ChR2-expressing ad libitum fed mice (Figure 3F).

**Figure 3.**
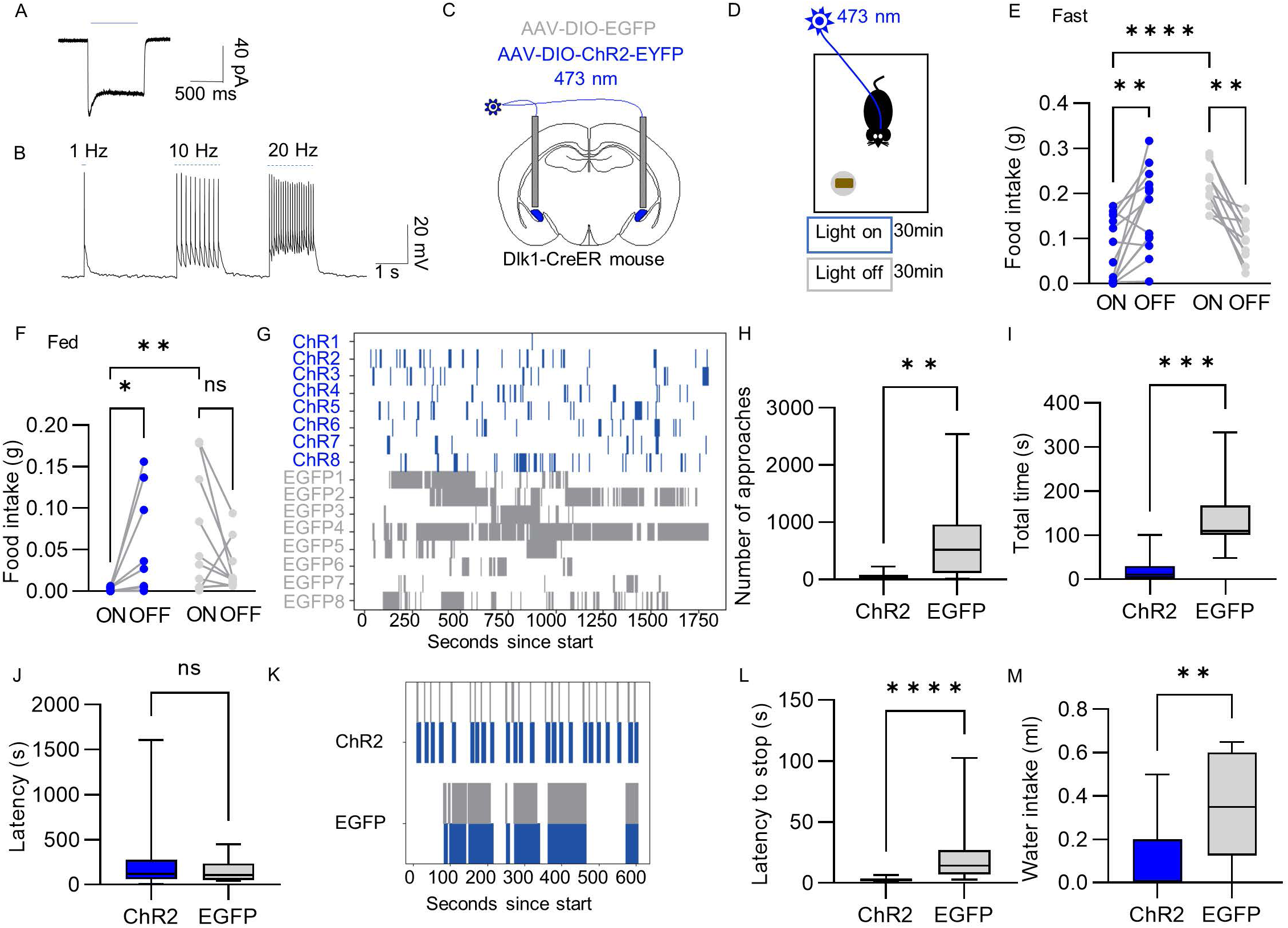
Activation of CeA^Dlk1^ neurons inhibits feeding. **A** An evoked current induced by photoactivation of CeA^Dlk1^ neurons. **B** Brain slice whole-cell patch-clamp recording traces showing action potentials triggered in CeA^Dlk1^ neurons expressing ChR2 by 1 Hz, 10 Hz, 20 Hz pulses. **C** Optic-fiber placement above Dlk1-CreER::ChR2-expressing or Dlk1-CreER::EGFP-expressing neurons. D Schematic of the paradigm for testing the effects of photoactivation on feeding behavior. **E** Food intake by fasted animals expressing control protein EGFP (n = 10) or ChR2 (n = 14). Blue is ChR2 group, and grey is EGFP. Data were analyzed by two-way ANOVA. For ChR2 versus EGFP light-ON: P < 0.0001. For ChR2 light-ON versus ChR2 light-OFF, P = 0.0023. For EGFP light-ON versus EGFP light-OFF, P = 0.0028. **F** Food intake by fed animals (same groups as in panel E). Data were analyzed by two-way ANOVA. For ChR2 versus EGFP light-ON: P = 0.0075. For ChR2 light-ON versus ChR2 light-OFF, P = 0.0428. ns, not significant. **G** Raster plot of food approaches of 20h fasted individual mice from the photoactivated ChR2 and EGFP groups. **H** Numbers of food approaches of 20h fasted photoactivated ChR2 and EGFP groups. Two-tailed unpaired t test, P = 0.0069. **I** Total time of food approach of 20h fasted photoactivated ChR2 and EGFP groups. Two-tailed unpaired t test, P = 0.0002. **J** Latency of the first approach to food of 20h fasted photoactivated ChR2 and EGFP groups. Two-tailed unpaired t test, P = 0.3839. **K** Raster plots showing feeding episodes recorded in the home cage by example animals expressing control protein EGFP or ChR2. Activation light (473 nm) was triggered 1–2 s after onset of feeding. 20Hz, 10-ms light pulses were delivered for 10 s. **L** Latency to stop feeding in response to photostimulation. Two-tailed unpaired t test, P < 0.0001. **M** Water intake by water deprived photoactivated ChR2 and EGFP groups. Two-tailed unpaired t test, P = 0.0096. Box–whisker plots display median, interquartile range and fifth to ninety-fifth percentiles of the distribution. *P < 0.05; **P < 0.01; ***P < 0.001; ****P < 0.0001.

To gain more insight into the inhibitory effects of CeA^Dlk1^ neurons on feeding, we performed a time-resolved analysis of feeding in mice that were deprived of food for 20 h. This analysis revealed that photoactivation of CeA^Dlk1^ neurons decreased the number of food approaches (defined as the time point at which the animal’s nose touched the food pellet) (Figure 3G, H), and decreased the total time of food contact (Figure 3I), whereas the latency to approach the food remained unchanged compared to photoactivated EGFP control mice (Figure 3J). Other parameters of locomotor activity such as total distance moved and velocity were not significantly different between ChR2-expressing mice and control mice (Figure S4B, C).

Next, we tried to obtain independent confirmation of our optogenetics results with a chemogenetic approach. We expressed Cre-dependent stimulatory hM3Dq designer receptors exclusively activated by designer drugs (DREADDs) selectively in CeA^Dlk1^ neurons and activated the neurons by intraperitoneal injection (i.p.) of the DREADD ligand clozapine-N-oxide (CNO). Chemoactivation of satiated Dlk1-CreER::hM3Dq mice resulted in decreased food intake in comparison with mice that received saline i.p. injections or with CNO-treated control mice expressing mCherry (Figure S4D). Then, we asked if photoactivation of CeA^Dlk1^ neurons was sufficient to interrupt ongoing feeding. The mice were placed in their home cages and a 10s photoactivation pulse was triggered 1∼2s after the onset of feeding. The results revealed that photoactivation of CeA^Dlk1^ neurons rapidly and consistently interrupted ongoing feeding within a few seconds following photostimulation (Figure 3K-L and Supplementary Movie A, B). Knowing that CeA^Dlk1^ neurons respond to bitter tasting liquid, we also asked if these neurons could suppress water intake. Indeed, in a free drinking assay, photoactivation of CeA^Dlk1^ neurons suppressed water intake by water-deprived mice (Figure 3M). In summary, these results indicate that activation of CeA^Dlk1^ neurons was sufficient to inhibit food and water intake.

### Activation of CeA^Dlk1^ neurons does not induce anxiety, but defensive behavior

Next, we asked if the observed inhibition of food intake by CeA^Dlk1^ neurons may in part be due to increased anxiety or defensive behavior. We photoactivated CeA^Dlk1^ neurons and tested the mice in open field (OF) and elevated plus maze tests (EPM) which assess anxiety-related behaviors. In the OF test, reduced time spent in the center zone of the arena is a measure of increased anxiety. Photoactivated Dlk1-CreER::ChR2 mice spent similar times in the center zone both in light ON and OFF phases, and compared to photoactivated Dlk1-CreER::EGFP control mice (Figure 4A,B). Other parameters such as frequency of entries into the center zone, total distance moved and velocity were not significantly different between Dlk1-CreER::ChR2 mice and control mice (Figure S5A-C).

**Figure 4.**
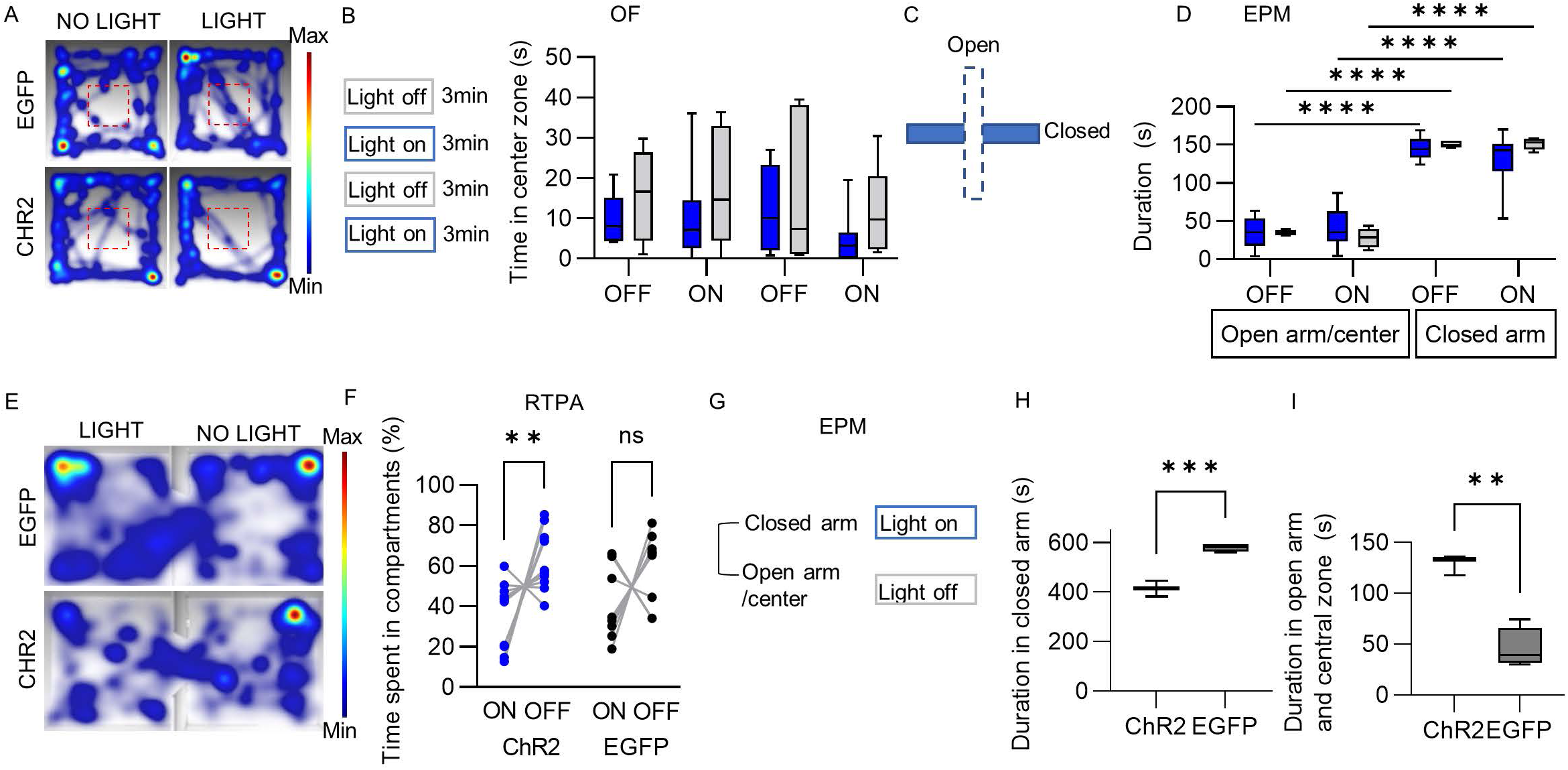
Activation of CeA^Dlk1^ neurons does not induce anxiety, but defensive behavior. **A** Representative images of heatmaps depicting the time spent at different locations in an open field box. The red box represents the center zone. **B** Photostimulation was set to the sequence (3min light-off, 3min light-on, 3min light-off, 3min light-on). Time spent in the central zone of the arena by animals expressing control protein EGFP (n=6) or ChR2 (n=10). Blue is ChR2 group, and grey is EGFP. Data were analyzed by Unpaired t test. P(OFF1) = 0.2102. P(ON1) = 0.2788. P(OFF2) = 0.6089. P(ON2) = 0.1141. **C** Schematic representation of elevated plus maze (EPM) with photostimulation delivered in the entire arena (same sequence in panel A). **D** Time spent in opens arms/central zone and closed arms by animals expressing control protein EGFP (n=6) or ChR2 (n=9). Data were analyzed by two-way ANOVA. ****P < 0.0001. **E** Representative images of heatmaps depicting the time spent in the RTPA task. **F** Preference of ChR2(n=10) and EGFP(n=9) mice for the photostimulation chamber. Data were analyzed by two-way ANOVA. For ChR2 light-On versus ChR2 light-Off, P = 0.0058. For EGFP light-On versus GFP light-Off, P = 0.1019. **G** A modified version of the EPM test. Light is turned on when mouse enters one of the two closed arms and turned off when leaving the closed arm. **H** Time spent in closed arms by animals expressing control protein EGFP or ChR2. Two-tailed unpaired t test, P=0.0003. **I** Time spent in opens arms and central zone by animals expressing control protein EGFP or ChR2. Two-tailed unpaired t test, P=0.0012.

In the EPM test, the innate response of mice is an aversion towards the exposed open arms and a center zone, resulting in an avoidance of these areas, and a preference for the sheltered space provided by the closed arms. We found that Dlk1-CreER::ChR2 and EGFP control mice spent most of the time in the closed arms. We also found that Dlk1-CreER::ChR2 mice spent similar times in the open arm/center of the arena during light ON phases than light OFF, and compared to photoactivated than Dlk1-CreER::EGFP control mice (Figure 4C,D).

To assess if CeA^Dlk1^ neuron activation elicits defensive behavior, we tested photoactivated Dlk1-CreER::ChR2 mice in the real-time place aversion paradigm (RTPA). Here, the mouse moves around freely in a two-compartment chamber, in which entry into one of two compartments is paired with photostimulation of CeA^Dlk1^ neurons. An aversive response to photostimulation would be displayed as avoidance of entry into the light-paired compartment, and hence avoidance of activation of CeA^Dlk1^ neurons. Photoactivated Dlk1-CreER::ChR2 mice spent significantly less time in the light-paired than the unpaired compartment showing an active avoidance of CeA^Dlk1^ neuron photostimulation. In contrast, control mice spent similar amounts of time in the two compartments (Figure 4E, F).

To investigate the strength of the aversive effect induced by CeA^Dlk1^ neuron photostimulation, we tested place aversion in a modified EPM setup, in which CeA^Dlk1^ neurons were activated directly upon entry into one of the two closed arms and turned off when leaving them, while entries into open arms or the center zone were not paired with CeA^Dlk1^ neuron stimulation (Figure 4G). Photoactivated Dlk1-CreER::ChR2 mice spent significantly less time in the closed arm and more time in the open arms and in the center compared to control mice, indicating active avoidance of the much preferred closed arms (Figure 4H,I). These results showed that the aversive effect induced by CeA^Dlk1^ neuron activation was stronger than the naturally aversive effect provided by unsheltered exposure. Together, these results indicate that CeA^Dlk1^ neuron activation does not promote anxiety, but real-time place aversion. Hence, the effects of CeA^Dlk1^ neuron activation on feeding may in part be due to modulating the feeding circuits and in part enhanced by increasing place aversion.

### Silencing CeA^Dlk1^ neurons overcomes the feeding block induced by nausea

Visceral malaise and bitter food are known to suppress appetite ^3^. Given that these conditions induced c-Fos expression in CeA^Dlk1^ neurons, we tested whether pharmacogenetic inhibition of CeA^Dlk1^ neuronal activity could overcome the effect of LiCl. For this purpose, we bilaterally injected the CeA of Dlk1-CreER mice with an AAV expressing the inhibitory DREADD (hM4Di), or mCherry (as a control) in a Cre-dependent manner (Figure 5A and S6A). Interestingly, chemoinhibition of CeA^Dlk1^ neurons with CNO rescued the effect of LiCl to inhibit feeding in food deprived animals to a level statistically indistinguishable from control mice that received saline instead of LiCl (Figure 5B). Administration of saline rather than CNO to LiCl-treated Dlk1-CreER::hM4Di mice failed to overcome the feeding block (Figure 5B). Similarly, CNO failed to rescue feeding in LiCl-treated Dlk1-CreER::EGFP control mice (Figure 5B).

**Figure 5.**
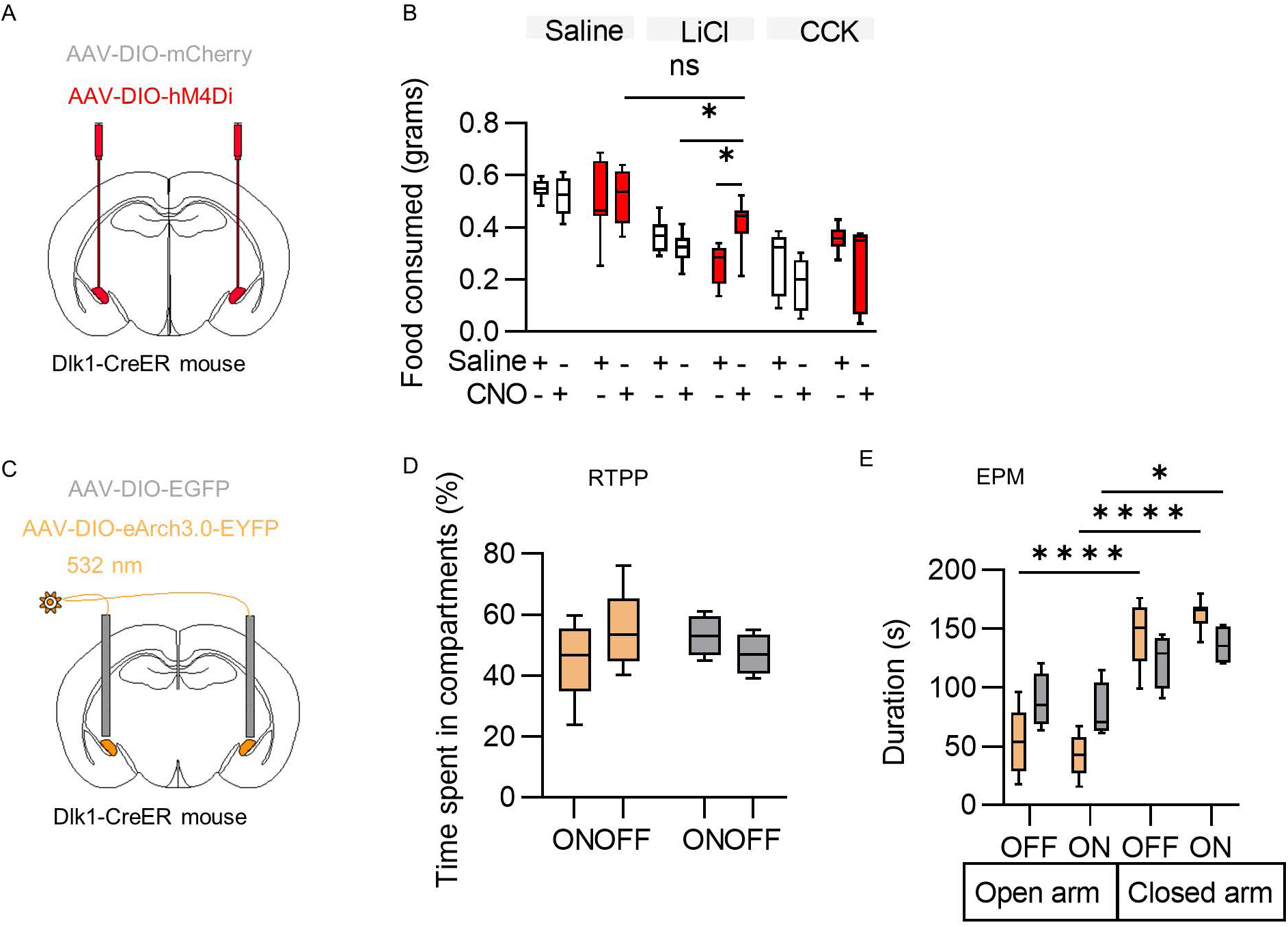
Silencing CeA^Dlk1^ neurons overcomes the feeding inhibition induced by nausea. **A** Delivery of AAV hM4Di-mCherry or AAV mCherry into the CeA in Dlk1-CreER mice. **B** Food intake in fasted animals after administration of different anorexigenic agent (n=6 mice per group). Data were analyzed by two-way ANOVA. Saline, F_(1,20)_ = 0.007, P = 0.9360; LiCl, F_(1,20)_ = 21.22, P = 0.0002; CCK, F_(1,20)_ = 0.016, P = 0.8994. **C** Optic-fiber placement above Dlk1-CreER::Arch-expressing or Dlk1-CreER::EGFP-expressing neurons. **D** Preference of Arch(n=9) and EGFP(n=4) mice for the photostimulation chamber. Data were analyzed by two-way ANOVA. For Arch light-On versus Arch light-Off, P = 0.3069. For EGFP light-On versus EGFP light-Off, P = 0.9662. **E** Time spent in opens arms/central zone and closed arm by animals expressing control protein EGFP (n=4) or Arch (n=9). Data were analyzed by two-way ANOVA. *P < 0.05; ****P < 0.0001.

Given that CCK, which mimics satiety, did not induce appreciable c-Fos expression in CeA^Dlk1^ neurons, we also tested if pharmacogenetic inhibition of CeA^Dlk1^ neurons could overcome the effect of CCK. Consistent with the c-Fos experiments, silencing of CeA^Dlk1^ neurons did not overcome the feeding block induced by CCK (Figure 5B).

Since photoactivation of CeA^Dlk1^ neurons had shown enhanced place aversion, we next asked, if photoinhibition would elicit real-time place preference (RTPP). As in the RTPA, in RTPP one of the two compartments is paired with light. To investigate this, we used a light-sensitive proton pump, archaerhodopsin (Arch) to inhibit CeA^Dlk1^ neurons in an optogenetic experiment. We first confirmed using whole-cell patch-clamp recordings in CeA brain slices from Dlk1-CreER mice injected with a Cre-dependent AAV expressing Arch, that 532-nm light pulses triggered photoinhibition in CeA^Dlk1^ neurons (Figure S6B-D). We bilaterally injected AAVs encoding Cre-inducible Arch (AAV9-EF1a-DIO-eArch3.0-EYFP), or control virus (AAV2-EF1a-DIO-EGFP) into the CeA of Dlk1-CreER mice and implanted optic fibers bilaterally over the CeA (Figure 5C and S6E). We found that Dlk1-CreER::Arch mice did not exhibit a significant preference for the photoinhibition-paired chamber (Figure 5D and S6F), suggesting that inhibition of CeA^Dlk1^ neurons was not intrinsically rewarding. Other parameters such as distance moved and velocity did not change either (Figure S6G, H). To test whether inhibition of CeA^Dlk1^ neurons was anxiolytic, we subjected photoinhibited Dlk1-CreER::Arch and EGFP control mice to EPM and OF tests. For the EPM test, we found that Dlk1-CreER::Arch and EGFP control mice spent most of the time in the closed arms, but no differences of time spent were found between light ON and light OFF phases, and between Dlk1-CreER::Arch and EGFP control mice (Figure 5E). In the OF test, similar as EGFP control mice, Dlk1-CreER::Arch mice spent similar time in central zone during light ON and light OFF phases (Figure S6I). Other parameters of locomotor activity such as total distance moved and velocity were not significantly different between Dlk1-CreER::Arch mice and control mice (Figure S6J, K). Together, these results showed that inhibition of CeA^Dlk1^ neurons is unlikely to have a rewarding effect or to alter locomotion. They further suggest that CeA^Dlk1^ neurons are part of a CeA microcircuit that suppresses feeding under conditions of nausea, independent of circuits regulating satiety or anxiety.

### CeA^Dlk1^ and CeA^Pkcδ^neurons form overlapping, but distinct networks

We next sought to understand the microcircuits in which CeA^Dlk1^ neurons are operating, in comparison to the related CeA^Pkcδ^ neurons. To map the monosynaptic inputs to CeA^Dlk1^ neurons, we injected a Cre-dependent monosynaptic retrograde rabies system in the CeA of Dlk1-CreER mice (Figure 6A, B). We found that CeA^Dlk1^ neurons received direct inputs (RFP^+^ cells) mainly from amygdala, hypothalamus and thalamus, but also from cortex, pallidum, mid-, and hindbrain (Figure 6C). Most notable brain regions were the lateral hypothalamic area (LH), insular cortex (IC), posterior thalamic nuclei (PO), parasubthalamic nucleus (PSTh), and PVT (Figure 6D, E, S7A-D). Among the top 20 inputs, several have been implicated in emotional behavior and feeding control, including the PO, which could potentially process multisensory information related to aversive stimuli and drive aversive behaviors ^31^. The PSTh has been shown to play a critical role in appetite suppression ^32^. The IC has been associated with sensory experience and emotional valence ^33^ and piriform (PIR) and entorhinal cortices (EC) link odor processing and perception ^34^.

**Figure 6.**
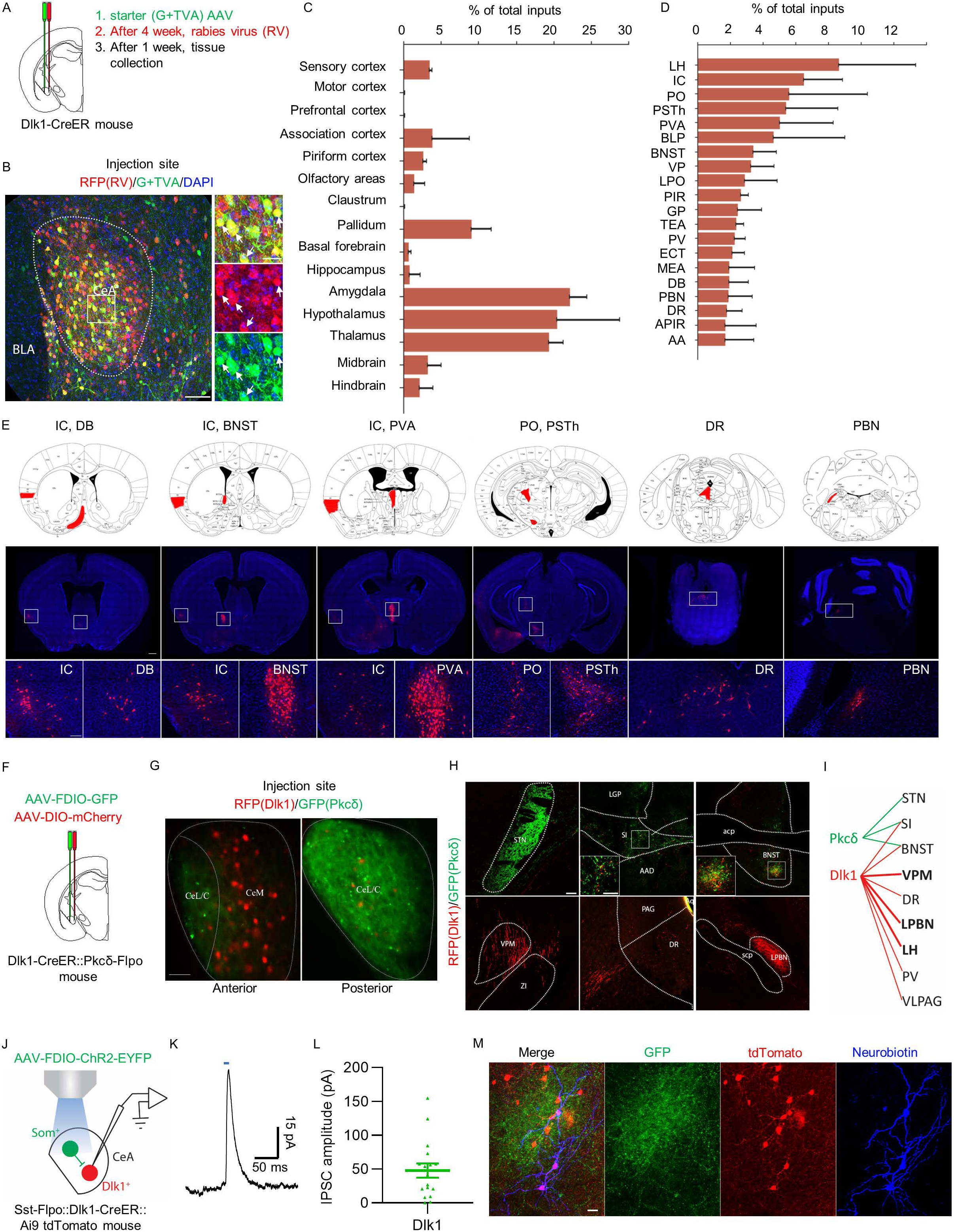
CeA^Dlk1^ neurons form a network regulating aversive behavior. **A** Schematic of the strategy for monosynaptic retrograde rabies virus (RV) tracing. **B** Representative image of the injection site. White box indicates the location of the high-magnification panel on the right. Arrows indicate starter cells expressing G + TVA (EGFP) and RV (RFP). Scale bar, 100 μm (left), 30 μm (right). **C** Major brain regions projecting to CeA^Dlk1^ neurons depicted as percentage of total inputs (n = 3 mice; error bars represent SEM). **D** Subregions projecting to CeA^Dlk1^ neurons (n = 3 mice; error bars represent SEM). Abbreviations see below. **E** Representative images of the areas projecting to CeA^Dlk1^ neurons. White box indicates the location of the high-magnification panel on the bottom. Scale bars, 500 μm (top), 100 μm (bottom). **F** Scheme of a Dlk1-CreER::Pkcδ-Flpo mouse injected into the CeA with AAVs expressing Cre-dependent mCherry and Flp-dependent GFP. **G** Representative images of the CeA showing mCherry and GFP expressing cells and their projections at anterior and posterior positions. Scale bar, 30 μm. **H** Representative images of brain areas innervated by CeA^Pkcδ^ neurons (green) and CeA^Dlk1^ neurons (red). White box indicates the location of the high-magnification panel. Scale bar, 100 μm. **I** Diagram illustrating brain regions innervated by CeA^Pkcδ^ neurons (green) and CeA^Dlk1^ neurons (red). Projections with large numbers of axons are indicated with thicker lines and bold font. **J** Scheme of whole-cell recording from CeA^Dlk1^ neurons in Sst-Flpo::Dlk1-CreER::Ai9 tdTomato slices transduced with Flp-dependent AAV-FDIO-ChR2-EYFP. **K** Light-evoked representative IPSCs in postsynaptic CeA^Dlk1^ neurons. Blue bar indicates the light. L Quantification of light-evoked responses (n=17 neurons; error bars represent SEM). **M** Representative image of ChR2-eYFP and tdTomato (Ai9) expression in the CeA. The recorded CeA^Dlk1^ neurons were neurobiotin-filled. Abbreviations: AA, anterior amygdaloid area; APIR, amygdalopiriform transition area; BNST, the bed nucleus of the stria terminalis; BLP, basolateral amygdaloid nucleus, posterior part; DR, dorsal raphe; DB, diagonal band; EA, temporal association cortex; ECT, Ectorhinal Cortex; GP, globus pallidus; IC, insular cortex; LH, lateral hypothalamic area; LPO, lateral preoptic nucleus; LPBN, lateral parabrachial nucleus; MEA, medial amygdaloid nucleus; PVA, paraventricular thalamic nucleus, anterior part; PO, posterior thalamic nuclear group; PSTh, parasubthalamic nucleus; PIR, piriform cortex; PV, paraventricular thalamic nucleus; PBN, Parabrachial nucleus; STN, subthalamic nucleus; SI, substantia innominate; VP, ventral pallidum; VPM, ventral posteromedial thalamic nucleus; VLPAG, ventrolateral periaqueductal gray.

Direct inputs of CeA^Pkcδ^ neurons have been previously mapped and shown to come from forebrain, cortex and hippocampus, followed by thalamus, basolateral amygdala (BLA), BNST, PSTh and PBN ^3, 21, 35^. Taken together, shared inputs between CeA^Pkcδ^ and CeA^Dlk1^ neurons include IC, PIR, BLA, BNST, PSTh, PBN and some parts of the thalamus. Interestingly, PBN cells that provide direct input to CeA^Dlk1^ neurons are not CGRP-positive (Figure S7E).

Next, we mapped the projection targets of CeA^Dlk1^ neurons in direct comparison with CeA^Pkcδ^ neurons. For this, we used Dlk1-CreER mice carrying in addition a newly generated Pkcδ-Flpo transgene (F. Fermani, P. Alcala, R.K., unpublished results). Pkcδ-Flpo expression faithfully represented endogenous Pkcδ expression (data not shown). Injection of a mixture of a Flp-dependent GFP reporter virus and a Cre-dependent mCherry reporter virus in the CeA of Dlk1-CreER::Pkcδ-Flpo mice, followed by Tamoxifen induction, revealed distinct red labeled CeA^Dlk1^ neurons and green labeled CeA^Pkcδ^ neurons and, as expected, very few double positive cells (Figure 6F,G).

We found that CeA^Pkcδ^ and CeA^Dlk1^ neurons had mostly unique projection targets and only few common targets. CeA^Dlk1^ neurons projected to a large number of unique mid – and hindbrain regions, including the ventral posteromedial nucleus of the thalamus (VPM), an important structure for conveying nociceptive information ^36^, the midbrain dorsal raphe nucleus (DR), a major source of neuromodulators, and responsible for the release of serotonin to other parts of the brain ^37^, and the lPBN, a hindbrain structure conveying interoceptive information ^20^ (Figure 6H, I and S7F). CeA^Pkcδ^ neurons showed one unique projection to the subthalamic nucleus (STN), which is associated with the inhibition of ongoing behavior and a critical hub for aversion ^38^. Both cell populations projected to SI and BNST, structures involved in negative reinforcement learning and negative emotion, respectively ^9^. These results were consistent with the model that CeA^Pkcδ^ neurons elicit anorexigenic behavior by inhibiting local appetitive neurons ^3^ and provide support for a model that CeA^Dlk1^ neurons function by inhibiting distant brain regions.

Next, we asked, how the activity of CeA^Dlk1^ neurons may be regulated. Given that CeA^Sst^ neurons were previously shown to locally inhibit CeA^Pkcδ^ neurons in the CeL ^39, 40^, we investigated if CeA^Sst^ neurons could inhibit CeA^Dlk1^ neurons which are enriched in the CeM. We generated triple transgenic Sst-FlpO::Dlk1-CreER::tdTomato mice and injected a Flp-dependent AAV-FDIO-ChR2 into the CeA. This allowed us to specifically photoactivate CeA^Sst^ neurons, while recording from tdTomato-expressing CeA^Dlk1^ neurons (Figure 6J). Whole-cell recordings of CeA^Dlk1^ neurons in brain slices revealed light-evoked, short latency, picrotoxin-sensitive IPSCs (17/17 neurons) (Figure 6K,L). Post hoc identification of neurobiotin-filled recorded neurons revealed that they were surrounded by the axons of CeA^Sst^ neurons (Figure 6M). These results suggest that the activities of anorexigenic CeA^Dlk1^ neurons are under inhibitory control of CeA^Sst^ neurons and raise the possibility that appetitive behaviors mediated by CeA^Sst^ neurons may in part be elicited or enhanced by inhibition of aversive CeA^Dlk1^ and CeA^Pkcδ^ neurons.

### The anorexigenic activity of CeA^Dlk1^ neurons is mediated by projections to the PBN

The PBN is mainly comprised of various glutamatergic neurons and scattered GABAergic neurons, that can inhibit local glutamatergic neurons ^41^. To elucidate which neurons in PBN receive CeA^Dlk1^ neuron input, we anatomically mapped the long-range projections of these neurons to the PBN. By selectively expressing a Cre-dependent EGFP (AAV-DIO-EGFP) in CeA^DLk^^1^ neurons (Figure 7A), we observed dense efferent fields most prominently in the lPBN (Figure 7B). Interestingly, we found a neuronal population expressing calretinin (Calb2), a calcium binding protein expressed by a subset of GABAergic neurons ^42^, tightly surrounded by the axons of CeA^Dlk1^ neurons (Figure 7B). Calb2^+^ neurons were located in the vicinity of CGRP neurons (Figure 7B, C, S8A) that had previously been shown to transmit threat signals to the extended amygdala and to inhibit feeding ^6^. This spatial arrangement of CeA^Dlk1^ projections – the potential inhibition of Calb2^+^ neurons and disinhibition of CGRP neurons (to inhibit feeding) – encouraged us to assess the functionality of this projection. We injected Cre-dependent ChR2-eYFP into the CeA in Dlk1-CreER mice (Figure 7D) and performed whole-cell recordings in brain slices from PBN neurons inside the area of ChR2 innervation (Figure 7E). We detected light-evoked IPSCs in most recorded PBN neurons (21/25) (Figure 7F, G). Post hoc identification of neurobiotin-filled neurons revealed that more than half of the recorded neurons were positive for Calb2 (Figure 7E).

**Figure 7.**
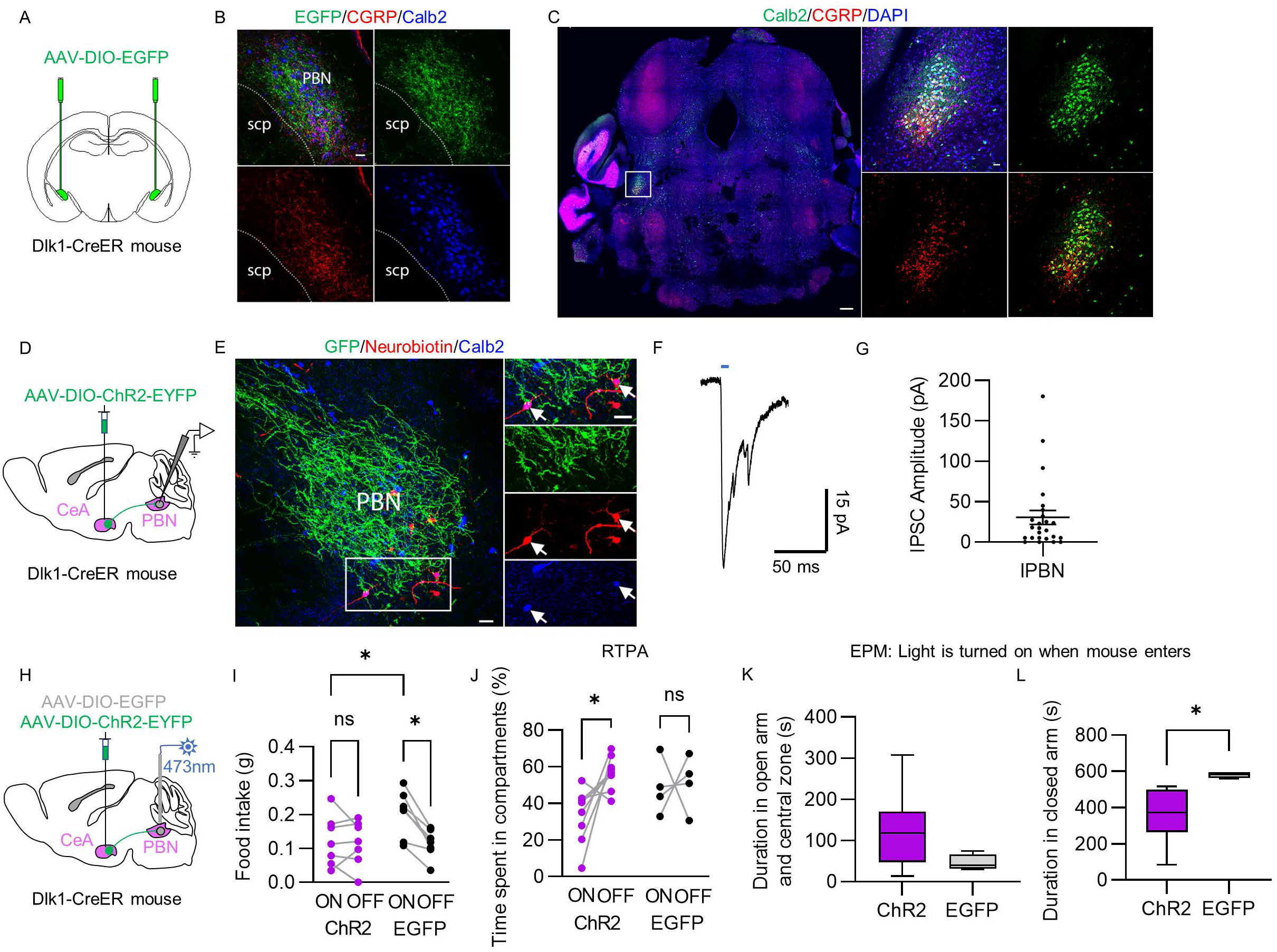
The anorexigenic activity of CeA^Dlk1^ neurons is mediated by projections to the PBN. **A** Delivery of AAV-DIO-EGFP into the CeA in Dlk1-CreER mice **B** ChR2-expressing axons of CeA^Dlk1^ neurons terminate in the lateral PBN with immunostaining for Calb2 (blue) and CGRP (red). Scale bar, 30um. **C** Representative image of PBN immunostaining for Calb2 (green) and CGRP (red). White box indicates the location of the high-magnification panel on the right. Scale bar, 100 um (left), 30um (right). **D** Targeting of AAV-DIO-ChR2-eYFP to the CeA^Dlk1^ neurons to investigate the Dlk1-CreER::ChR2→PBN projection. **E** Representative image of ChR2-expressing axons of CeA^Dlk1^ neurons terminating in the lateral PBN. Recorded neurons were neurobiotin-filled (red), immunostained with Calb2 (blue). White box indicates the location of the high-magnification panel on the right. Arrows indicate neurobiotin and Calb2 double positive cells. Scale bar, 30um. **F** Light-evoked representative IPSCs in postsynaptic PBN neurons. Blue bar indicates the 3 ms of light stimulation. **G** Quantification of light-evoked responses (n = 25 neurons; error bars represent SEM). **H** Delivery of AAV-DIO-ChR2-eYFP or AAV-DIO-EGFP bilaterally into the CeA with a bilateral fiber-optic implant over the PBN (only one side is shown). **I** Food intake by Dlk1-CreER::ChR2 → PBN (n=7, magenta) and control mice (n=7, black) when optogenetically stimulating the CeA^Dlk1^ terminals in PBN. Data were analyzed by two-way ANOVA. For ChR2 light-On versus EGFP light-On, P = 0.0268. For EGFP light-On versus EGFP light-Off, P = 0.0151. **J** Time spent by Dlk1-CreER::ChR2 → PBN (n=8) and control mice (n=4) in the photostimulated side of the RTPA chamber. Data were analyzed by two-way ANOVA. For ChR2 light-On versus ChR2 light-Off, P = 0.0137. **K** Time spent in open arm and central zone by Dlk1-CreER::ChR2 → PBN (n=9) and control mice (n=4). Light is turned on when mouse enters closed arm. Two-tailed unpaired t test, P = 0.1389. **L** Time spent in closed arm by Dlk1-CreER::ChR2 → PBN (n=9) and control mice (n=4). Light is turned on when mouse enters closed arm. Two-tailed unpaired t test, P = 0.0129.

Next, we asked whether photoactivation of CeA^Dlk1^ axons projecting into the lPBN would be sufficient to mediate feeding suppression. We transduced CeA^Dlk1^ neurons with Cre-dependent ChR2-eYFP and placed optic fibers bilaterally above the PBN (Figure 7H). In a 30 min light-ON followed by 30 min light-OFF feeding paradigm, photoactivation of the CeA^DLk^^1^-neuron presynaptic terminals in the PBN elicited a significant decrease in food intake in fasted Dlk1-CreER::ChR2→PBN compared to photoactivated Dlk1-CreER::EGFP→PBN control mice. (Figure 7I). In the following 30 min light-OFF phase, control mice consumed significantly less food, because they were satiated. In contrast, Dlk1-CreER::ChR2→PBN mice were only partially satiated and showed similar feeding behavior as in the light-ON phase. The time-resolved analysis of feeding in Dlk1-CreER::ChR2→PBN and Dlk1-CreER::EGFP→PBN mice revealed that photoactivation of CeA^Dlk1^ presynaptic terminals in the PBN decreased the number of food approaches, and the total time of food contact, whereas the latency to approach the food remained unchanged compared to photoactivated Dlk1-CreER::EGFP→PBN mice (Figure S8B-E). We also probed whether activation of the CeA^Dlk1^ neuron−PBN projection had an aversive effect. Indeed, 20-Hz photoactivation of Dlk1-CreER::ChR2→PBN mice elicited real-time place avoidance without changing locomotor activity such as total distance moved and velocity (Figure 7J and S8F-H). It also reduced the natural preference for a sheltered environment in the EPM compared to photoactivated Dlk1-CreER::EGFP→PBN control mice (Figure 7K, L). Together, our results indicate that CeA^Dlk1^ projections to the PBN are sufficient to inhibit feeding and to drive place avoidance behavior, possibly through inhibition of a population of PBN GABAergic neurons marked by expression of Calb2.

## Discussion

Here, we characterized CeA^Dlk1^ neurons as a newly identified CeA population enriched in the medial subdivision (CeM). We found that CeA^Dlk1^ neurons are activated by both LiCl, which causes nausea, and quinine, which is a bitter tastant. Photoactivation of these neurons inhibits feeding and drinking behavior and has an aversive effect, which does not appear to be due to anxiety or altered locomotor activity. Conversely, photoinhibition of CeA^Dlk1^ neurons overcomes the anorexigenic effect induced by LiCl. Hence, CeA^Dlk1^ neurons are the first CeM neurons with aversive and anorexigenic properties and they are distinct from aversive CeA^Pkcδ^ neurons that reside in the CeL. Whereas CeA^Pkcδ^ neurons are activated by satiety signals and nausea, CeA^Dlk1^ neurons are specifically activated by nausea, but not satiety.

The CeA has been a focus of investigation in recent scRNA-Seq and mFISH studies, leading to the discovery of several new cell types and their spatial locations. CeA^Dlk1^ neurons were characterized as a novel transcriptomic cell cluster located in the CeM and interphase between CeL/CeM ^14, 22^. In a separate study ^15^, a cell cluster with similar transcriptome was characterized (named Nr2f2 cluster) and found to have the same spatial location. We find that the Nr2f2 cluster consists mostly of Dlk1 neurons and some contribution by PKCδ neurons (W.D., unpublished observations). CeA^Dlk1^ neurons are therefore the third CeA population with predominantly aversive functions, each being located in a different subdivision: CeA^Calcrl^ neurons in CeC, CeA^Pkcδ^ neurons in CeL, and CeA^Dlk1^ neurons in CeM. Our ex vivo recordings revealed that the physiological properties of CeA^Dlk1^ neurons are more similar to CeA^Pkcδ^ neurons, than to predominantly appetitive CeA^Sst^ neurons. They have a higher membrane potential than CeA^Sst^ neurons, suggesting CeA^Dlk1^ neurons may require a stronger depolarizing input to reach the threshold for firing an action potential.

The most distinctive aspect of CeA^Dlk1^ neurons is their rich and diverse projection output. Whereas CeA^Pkcδ^ neurons mostly project to targets within the CeA and few long-range targets outside the CeA, such as BNST and SI, CeA^Dlk1^ neurons form efferent projections to LH, VPM, lPBN, DR, PVT, vlPAG, BNST and SI. CeA^Pkcδ^ neurons likely exert their anorexigenic functions by inhibition of local appetitive CeA neurons ^3^, thereby making the food less palatable. Instead, CeA^Dlk1^ neurons mediate their aversive effects at least in part through inhibition of neurons in the PBN and this projection likely has a direct effect on feeding by either inhibiting appetitive input to the PBN or disinhibiting aversive PBN neurons (Figure 8). Candidate target neurons for the latter mechanism are inhibitory Calb2^+^ PBN neurons whose activity is inhibited by CeA^Dlk1^ projections. This effect may indirectly disinhibit aversive CGRP^+^ PBN neurons and promote suppression of feeding. Alternatively, inhibitory CeA^Dlk1^ projections to the PBN may interfere with the activity of inhibitory appetitive CeA projections coming from CeA^Htr2a/Pnoc^ or CeL^NTS^ neurons, for example through a presynaptic inhibition mechanism. These putative circuit mechanisms remain to be demonstrated. It also remains to be determined, if CeA^Dlk1^ neurons locally inhibit appetitive CeM neurons and thereby reduce appetite.

**Figure 8.**
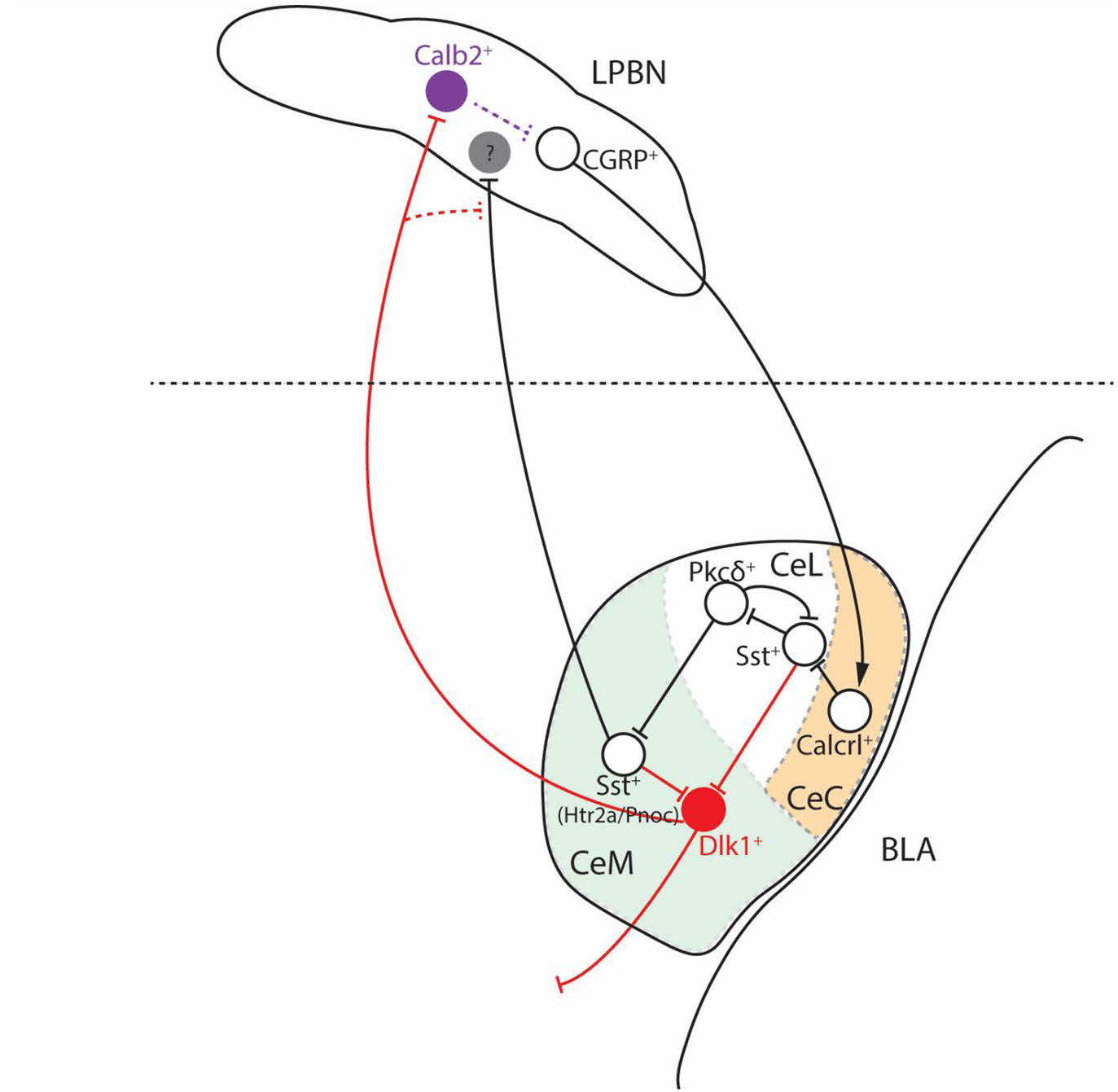
Schematic model of the CeA^Dlk1^ microcircuit. CeA^Dlk1^ neurons are predominantly located in CeM, while the other anorexigenic neurons are in CeL (PKCδ) and in CeC (Calcrl). CeA^Dlk1^ neuron activity is likely regulated by appetitive CeA neurons, such as Sst^+^ neurons in CeL and CeM, and by long-range inputs (not shown here). Activation of the CeA^Dlk1^→PBN projection is sufficient to suppress feeding, possibly by inhibition of Calb2^+^ neurons in lPBN. Whether their inhibition disinhibits nearby CGRP^+^ neurons remains to be shown. CGRP^+^ neurons in lPBN activate Calcrl^+^ neurons in CeC ^6, 7^. CeL^PKCδ^ neurons exert their activity mainly by inhibiting local appetitive CeA neurons ^3^. Appetitive CeM neurons such as Htr2a^+^ and Pnoc^+^ neurons, also project to the PBN to promote feeding ^21–23^. Their target cells in PBN are unknown. CeA^Dlk1^→PBN projectors may inhibit the appetitive projections by presynaptic inhibition. The dashed line indicates CeA and lPBN are located at different bregma levels. **Movie** Light activation of CeA^Dlk1^ neurons inhibits feeding. **A** This video shows a 20 h fasted mouse that approaches and eats a food pellet in its home cage, its feeding behavior after photostimulation of CeA^Dlk1^ neurons that express ChR2. **B** This video shows a 20 h fasted mouse that approaches and eats a food pellet in its home cage, its feeding behavior after photostimulation of CeA^Dlk1^ neurons that express control EGFP.

Activation of CeA^Dlk1^ neurons also produced avoidance behavior in a real-time place aversion assay, but did not alter anxiety-related behaviors in open field or elevated plus maze tests. Some of the projection targets of CeA^Dlk1^ neurons have reportedly critical roles in avoidance behavior and feeding behaviors. For example, CeA projections to LH are preferentially activated in male rats that show avoidance of a predator odor-paired context. LH Hcrt^+^ cells, which receive input from CeA, become activated during predator odor exposure and optical stimulation of LH Hcrt^+^ cells produces real-time place avoidance ^43, 44^. Hence, CeA^Dlk1^ neuron projections to LH may be an interesting circuit for future investigations.

Other interesting aspects of CeA^Dlk1^ neurons are their activation mechanisms and afferent inputs. We found CeA^Dlk1^ neurons to be activated by nausea and bitter tastants, but, unlike CeA^Pkcδ^ neurons, not by satiety signals, suggesting that their role in feeding behavior may be more specific towards danger signals associated with unpalatable or poisonous food. Previous studies have established the neuronal circuitry through which CeA^Pkcδ^ neurons mediate the effects of satiety ^45^, while the CeA circuit that mediates the effect of nausea is largely unknown. Local connectivity and functional properties of projection neurons in amygdala can depend on their long-range targets ^46^. Our present data suggests that CeA^Pkcδ^ and CeA^Dlk1^ neurons make complementary contributions to anorexigenic behavior.

Electrophysiology data suggests that CeA^Dlk1^ neurons receive more inhibitory inputs and fewer excitatory inputs than Sst^+^ neurons. Inhibitory inputs may come from local Sst^+^ neurons or from distant brain regions. We have shown that CeA^Sst^ neurons locally inhibit CeA^Dlk1^ neurons. Our method did not allow us to distinguish whether the inhibition derived from CeA^Sst^ neurons in CeL, CeM, or both. It remains to be determined whether the inhibitory control happens within the same subdomain (CeM to CeM) or across subdomain (CeL to CeM).

Rabies tracing indicated that CeA^Dlk1^ neurons receive monosynaptic inputs from multiple input regions including hypothalamus, thalamus, and insular cortex. Long-range inhibitory inputs may come from BNST and DR. Previous work showed that BNST to CeA projections predominantly arise from GABAergic neurons ^47^. CeA^Dlk1^ neurons also project to BNST. Antagonistic interactions between BNST and CeA likely determine the intensity and specificity of aversive responses in threatening conditions ^48^. Serotonergic neurons that co-express corticotropin-releasing factor (CRF) in DR give inhibitory projections to CeA. DR is a major source of serotonin, and the dopamine system receives common inputs from brain regions associated with appetitive and aversive information processing ^49^.

PVT was one of the important excitatory inputs of CeA^Dlk1^ neurons. It has been demonstrated that the PVT-CeA circuit regulates a variety of behaviors, such as fear learning ^50–52^, reward-seeking ^53, 54^, anxiety ^55^, depression-related behavior ^56^, neuropathic pain ^57^, wakefulness control ^58^, and stress-sensitive repetitive behaviors ^59–61^. PVT integrates and amplifies cortical signals related to threats and relays these signals to activate subcortical circuits that modulate defensive responses including avoidance of potential threats ^62^.

PBN projections to the CeA have been proposed to serve as a general alarm system to potential threats. CeA^Pkcδ^ neurons in the CeC region, labelled by Calcrl, are involved in suppressing food intake by receiving projections from calcitonin gene related peptide (CGRP) neurons in the lPBN, which transmit anorexigenic and danger signals to the CeA ^6, 7^. CeA^Dlk1^ neurons also receive direct inputs from the PBN, but these were not among the strongest inputs. Moreover, PBN cells that provide direct input to CeA^Dlk1^ neurons are not CGRP-positive. In general, CeA^Dlk1^ neurons receive monosynaptic inputs from many brain areas implicated in feeding, energy balance, aversive information processing and taste and odor perception. This is consistent with previous studies demonstrating that the CeA integrates sensory and physiological information to regulate feeding and energy balance.

In conclusion, the present manuscript provides a detailed characterization of a newly identified CeA population that is important for nausea-induced food suppression. Our findings indicate that the anorexigenic effects of nausea, bitter tastants and satiety may be regulated by different neurons and complementary circuits in the CeA. In the future, further characterization of these circuits, including modulation by neuropeptides, in humans or non-human primates may provide insights into the etiology of obesity, anorexia or other eating disorders and may help to develop therapeutic strategies targeting these common and burdensome disorders.

## Acknowledgements

We thank Patrick Hoffmann, Ekaterina Prilevskaya and Fady Shenouda for help with management of the animal colony; Wenkang Wang (Max Plank Institute of Intelligent Systems) for help with data analysis; Karl-Klaus Conzelmann (Gene Center Munich, LMU) for providing EnvA G-deleted rabies virus. This study was supported by the Max-Planck Society and the European Research Council under the European Union’s Horizon 2020 research and innovation programme (no. 885192, BrainRedesign).

## Author contributions

WD and RK conceptualized and designed the study. WD conducted the experiments and analyzed data. HW assisted with behavior and histology experiments. CP performed electrophysiology experiments and assisted with mouse surgery. WD and RK wrote the paper with input from all authors. R.K. supervised and provided funding.

## Methods

### Animals

Male and female mice that were at least 2 months old were used in all experiments, following regulations from the government of Upper Bavaria. The mice were housed in their home cages with 2-6 mice per cage, under a 12-hour light/12-hour dark cycle, with food and water freely available. The Dlk1-CreER transgenic line (Tg(Dlk1-cre/ERT2)26.10Ics) were purchased from CERBM-GIE, PHENOMIN-Mouse Clinical Institute. Sst-Flp mice (stock number 031629) were purchased from Jackson Laboratories. The Td-Tomato (B6.Cg-Gt(ROSA)26Sor^tm9(CAG–tdTomato)Hze/J)63^ and Rosa26R^64^ mouse lines were as previously described. The Htr2a-Cre BAC transgenic line (stock Tg(Htr2a-Cre)KM208Gsat/Mmucd) BAC mice were imported from the Mutant Mouse Regional Resource Center (https://www.mmrrc.org/). Prkcd-Flpo transgenic mice were generated by BAC injection. The original BAC RP23-283B12 was modified by homologous recombination in bacteria to insert the codon optimized version of the Flpe site specific recombinase (Flpo) under the Prkcd (protein kinase C delta) regulatory sequences. Transgenic mice were bred onto a C57BL/6N background.

For optogenetic and chemogenetic manipulations, animals were handled and housed singly on a 12-hour inverted light cycle for at least 3 days prior to the experiments. Except during food deprivation for feeding experiments, mice were given ad libitum food access. All feeding behavior tests were carried out at the same time each day during the dark period (1 p.m.–6 p.m.).

### Viral vectors

The following AAV viruses were obtained from Addgene: AAV9-pAAV-hSyn-DIO-hM3D(Gq)-mCherry (Addgene, 44361), AAV9-pAAV-hSyn-DIO-hM4D(Gi)-mCherry (Addgene, 44362), AAV2-pAAV-hSyn-DIO-mCherry (Addgene, 50459), AAV5-Ef1a-DIO-ChR2-EYFP (Addgene, 35509), AAV5-Ef1a-DIO-EGFP (Addgene, 27056). AAV5-EF1a-DIO-eArch3.0-EYFP, AAV5-EF1a-fDIO-hChR2(H134R)-EYFP-WPRE, and AAV-synP-DIO-STPEPB were produced at the Gene Therapy Center Vector Core at the University of North Carolina Chapel Hill. EnvA G-deleted rabies for long-range monosynaptic tracing was a gift from Karl-Klaus Conzelmann (Gene Center Munich, LMU).

### snRNA-seq analysis

The snRNA-seq data was downloaded from public repository GEO (GSE231790). The t-distributed stochastic neighbor embedding (t-SNE), differential expression analysis and cell-type taxonomy tree construction were performed using Seurat V4.0.1.

### Stereotactic surgery

Mice were anesthetized with 1.5-2% isoflurane and received oxygen at 1.0 liter per minute before being placed in a stereotaxic frame (Kopf Instruments). A heating pad was used to keep the body temperature stable. Carprofen (5 mg/kg bodyweight) was used subcutaneously as an analgesic.

Once the mouse skull was exposed, we drilled a cranial window (1–2 mm^2^) unilaterally (in vivo photometry, monosynaptic rabies and anterograde tracing experiments) or bilaterally (optogenetic and chemogenetic experiments). Next, a glass pipette (#708707, BLAUBRAND intraMARK) was lowered into the window to deliver 300 nL of viral vector to the area of interest (coordinates: CeA: −1.22 mm anterior to bregma, ± 2.8 mm lateral from midline and, −4.7 mm vertical from the brain surface; PBN: −5.1 mm anterior to bregma, ± 1.7 mm lateral from midline and, −3.3 mm vertical from the brain surface). In the same surgery, mice used in optogenetic experiments were bilaterally implanted with optic fibers (200 μm core, 0.39 NA, 1.25-mm ferrule (Thorlabs)) above the CeA (−4.2 mm ventral) or PBN (−3.0 mm ventral). Implants were secured with cyanoacrylic glue, and the exposed skull was covered with dental acrylic (Paladur). For all other mice, the incision was closed with sutures.

Dlk1-CreER mice used for rabies tracing were first unilaterally injected in the CeA with the starter AAV-synP-DIO-STPEPB. After 4 weeks, the same mice were injected with the rabies virus. Seven days later, mice were killed, and their brain tissue was collected and processed for immunohistochemistry.

### Tamoxifen protocol

Tamoxifen solution was prepared in 90% corn oil and 10% ethanol. The injection dose was determined by weight (using approximately 200 mg tamoxifen/kg body weight) and was given for 5 consecutive days. For Dlk1-CreER::Ai9 tdTomato mice, we waited 1 week and then sacrificed the animals. For all AAV expression experiments, tamoxifen was administered 1 week after stereotactical AAV injection.

### Pharmacological treatments

For chemogenetic experiments, mice were given an intraperitoneal (IP) injection of CNO (0.4 mg/kg or 2 mg/kg diluted in saline) or the equivalent volume of saline before the experiment and were allowed to recover in their home cages for 30 minutes. For anorexigenic-drug studies, compounds used for intraperitoneal injection: LiCl (150 mg/kg) (Sigma) or CCK (5 μg/kg) (BACHEM). Behavioral tests were usually performed 20-30 min after drug injection. All drug treatments were administered in a counterbalanced fashion, with 2 days between experiments.

### Optogenetic manipulations

Mice were bilaterally tethered to optic fiber patch cords (Prizmatix) connected to a multi wavelength LED (Prizmatix) via a rotary joint (Prizmatix) and mating sleeve (Thorlabs). For photoactivation experiments, 10-ms, 473-nm light pulses at 20 Hz and 10–15 mW were used. Constant 532-nm light at 10 mW was used for photoinhibition experiments. The LED were triggered, and pulses were controlled with EthoVision XT16 software (Noldus Information Technologies).

### Feeding behavior

Mice were habituated to the behavioral context for daily 10-min sessions, for 2 days before the experiment. For 20h-fasted feeding test, mice were food restricted the day before test. Mice were presented with a regular food pellet, and allowed to feed. The weight of the food pellet, including the food debris left in the cage floor after test, was measured to calculate the food intake. For ‘fed’ feeding test, mice were not food deprived before the test. In the optogenetic experiment, the light was started just after the mice were put into the testing cage for 30 minutes, then the light was off for 30 minutes. The food intake was measured for both periods. For the pharmacological experiment, CNO or other compounds (LiCl and CCK) were injected 20-30 minutes before the test. The food intake was measured for 1h. All of the feeding tests were performed between 1 pm to 6 pm.

### Drinking behavior

The mice were habituated to the testing environment for 2 days with daily 10-minutes sessions before the experiment. Mice were water deprived for 20 hours the day before test. On the testing day, water deprived mice were given access to water, For the optogenetic experiment, the light was turned on for 30 minutes immediately after the mice were placed into the testing cage, then the light was turned off for 30 minutes. Water intake was measured during both periods. All of the drinking tests were performed between 1 pm to 6 pm.

### Real-time place preference/aversion test

Mice with optic fiber patch cables tethered were allowed to explore a two-compartment arena (50 cm × 25 cm × 25 cm). Mice were tested across two sessions. In session one, one side was assigned as the photostimulation chamber. Every time the mouse entered this chamber, 10-ms, 473-nm light pulses at 20 Hz and 10–15 mW (measured at the tip of optic fibers) were delivered intracranially for activation experiment, while constant 532-nm light at 10 mW were used for photoinhibition experiment. Photostimulation ceased when the mouse left the photostimulation side. In the second session, we assigned the other chamber as the photostimulation side and repeated testing. The behavior of the mice was recorded using a camera. Ethovision XT16 software (Noldus Information Technologies) was used to deliver light pulses and analyze behavioral parameters.

### Open-field (OF) test

Mice with optic fiber patch cables were tethered and allowed to explore a square arena (50 cm × 50 cm × 50 cm) for 12 minutes. Three minutes after mice were introduced to the arena, the photostimulation was delivered for 3 minutes, followed by 3 minutes without light, and then another 3 minutes with light. The spatial location and the movement of the mice were recorded and analyzed using video tracking software Ethovision XT16 (Noldus Information Technologies).

### Elevated plus maze (EPM) test

The EPM analysis was carried out in an apparatus with the shape of a plus, made of two open arms (35 cm length) and two closed arms (35 cm length) with walls (15 cm high), 5 cm wide and extended from a central platform (5 × 5 cm) to allow mice to freely move across the arms of the setup. The maze is elevated 65 cm from the floor. At the beginning of the 10min session, mice with optic fiber patch cables tethered were placed in the central zone with their heads towards the open arm. The photostimulation strategy was the same as in the OF test. The behavior of the mice was videotaped with a camera. Ethovision XT16 software (Noldus Information Technologies) was used to deliver light pulses and analyze behavioral parameters.

### Modified version of the EPM test

The setup is the same as the normal EPM test. The difference is in the timing of photostimulation. Photostimulation was activated upon entry into any of the closed arms, and turned off upon leaving them. In contrast, visiting the open arms or occupying the center zone had no effect on photostimulation. The animal was placed in the center zone of the arena and allowed to freely explore the apparatus for 10 minutes. The time spent in each compartment were recorded by the Ethovision XT16 tracking software (Noldus Information Technology).

### Long-range mapping of monosynaptic inputs

The mouse brains were selected based on high tracing efficiency and the presence of a large number of starter cells mainly restricted to the CeA. All images processing was performed using ImageJ software (NIH). For quantifications within all subregions, every section was analyzed using previously described methods ^65^, but only the input neurons ipsilateral to the injection site were counted. The central amygdala nucleus, as the input region, was excluded from the analysis. The numbers of input neurons for each experiment was normalized to the total number of inputs in each animal. Areas that contained less than 5% of the total inputs were excluded.

### Acute brain slice preparation and electrophysiological recordings

The mice were anesthetized using isoflurane. After ensuring a deep level of anesthesia, the animals were subjected to decapitation.

The freshly harvested brain was immediately immersed in an ice-cold cutting solution composed of the following components (mM): NaCl (30), KCl (4.5), MgCl2 (1), NaHCO3 (26), NaH2PO4 (1.2), glucose (10), and sucrose (194). This solution was equilibrated with a mixture of 95% O2 and 5% CO2, and the brain was then sectioned into slices of 280 μm thickness using a Leica VT1000S vibratome. The slices were subsequently transferred to an artificial cerebrospinal fluid (aCSF) solution, which contained (mM): NaCl (124), KCl (4.5), MgCl2 (1), NaHCO3 (26), NaH2PO4 (1.2), glucose (10), and CaCl2 (2). The aCSF solution was equilibrated with 95% O2/5% CO2 and maintained at a temperature of 30-32°C for 1 hour before being returned to room temperature.

The brain slices were mounted in a recording chamber and continuously perfused with the aforementioned aCSF solution equilibrated with 95% O2/5% CO2 at 30-32°C. Whole-cell patch-clamp recordings were performed using the techniques previously described in literature ^66^. The patch pipettes were prepared from filament-containing borosilicate micropipettes with a resistance of 5-7 MΩ. The intracellular solution used for recordings consisted of the following components (mM): potassium gluconate (130), KCl (10), MgCl2 (2), HEPES (10), Na-ATP (2), Na2GTP (0.2) and had an osmolarity of 290 mOsm. For PBN recordings in voltage-clamp we used the following intracellular solution: 125 mM CsCl, 5 mM NaCl, 10 mM HEPES, 0.6 mM EGTA, 4 mM Mg-ATP, 0.3 mM Na2GTP, 10 mM lidocaine N-ethyl bromide (QX-314), pH7.2, and 290 mOsm. The slices were visualized using an IR-DIC equipped fluorescence microscope (Olympus BX51) and data was obtained using a MultiClamp 700B amplifier, a Digidata 1550 digitizer, and analyzed using the Clampex 10.3 and Clampfit software from Molecular Devices. The data was sampled at 10 kHz, filtered at 2 kHz.

For optogenetic studies, stimulation of neurons was achieved using a multi-LED array system (CoolLED) connected to the aforementioned Olympus BX51 microscope.

### c-Fos activation experiment

Mice were given CCK (5 μg/kg) or LiCl (150 mg/kg) i.p. injections, while control mice received saline i.p. injection. Mice were sacrificed 90 min after the injection and tissue sections from CeA immunostained for c-Fos. For c-Fos expression analysis in fasted mice, food was removed for 20 h, afterwards the mice were sacrificed. Ad libitum fed mice were sacrificed during the dark (active) phase. Quinine water solution (10mM, 0.2ml) or plain water was gently applied (in 15sec) intraorally to normally hydrated mice using a standard ball-tipped gavage needle. Mice were perfused 90min after infusion.

### RNAscope experiment

Multicolor Fish was performed on fixed frozen brain slices. Briefly, Brain tissue was embedded in O.C.T. (Sakura Finetek, 4583) in cryomolds (Sakura Finetek, 4566) and fresh-frozen on dry ice. Brain tissue was cut on a cryostat (Leica, CM3050S) to obtain 20-μm sections, collected on slides (VWR Microslides Superfrost Plus, 48311-703) and stored at –80 °C. Hybridization was performed using the RNAscope kit (ACDBio). The protocol was previously described^67^. Probes against Dlk1 (405971-C2), LacZ (313451-C3), Nts (420441), Sst (404631), Drd2 (406501), Tac2 (446391), Pnoc (437881), Crh (316091-C3), and Prkcd (441791-C3) were applied at a 1:50 dilution to sections. Images were taken using a Leica SP8 confocal microscope and processed using ImageJ software (NIH).

### Immunohistochemistry

Brain slices were fixed in 4% paraformaldehyde in PBS, pH 7.4, permeabilized with 0.1% Triton X-100 in PBS for 15 min, and blocked with 10% donkey serum in PBS for 2 h at room temperature (RT). Antibody incubations were performed overnight at 4°C, followed by three washes in PBS. Fluorescent-conjugated secondary antibody incubation for 2h at RT. Images were acquired using a Leica SP8 confocal microscope with ImageJ software (NIH) and Imaris (Bitplane). Primary antibodies used were as follows: LacZ (1:200, ab9361, Abcam), GFP (1:500, A10262, Thermo Fischer Scientific, USA), cfos (1:750, 2250S, Cell Signaling, USA), CGRP (1:500, ab36001, Abcam), Calb2 (1:500, CR7697, swant). Secondary antibodies used were as follows: donkey anti-rabbit/goat/chicken Alexa Fluor 488/Cy3/Alexa Fluor 647 (1:500) (Jackson; anti-rabbit, 711-545-152, 711-165-152, 711-495-152; anti-chicken, 703-545-155, 703-165-155, 703-605-155). DNA was stained with 49,6diamidino-2-phenylindole (1:10000, D1306; Invitrogen).

### Statistical analysis

All statistics are described where used. Statistical analyses were conducted using GraphPad Prism 9 software (GraphPad). No statistical methods were used to predetermine sample sizes, but the number of samples in each group were similar to those reported in previous publications. Pairwise comparisons were calculated with unpaired or paired two-tailed t-tests, and multiple group data comparisons were calculated with two-way ANOVA with Bonferroni post hoc test. Data collection and analysis were not performed blind to the conditions of the experiments. Mice that, after histological inspection, had the location of the viral injection (reporter protein) or of the optic fiber(s) outside the area of interest were excluded. All data were represented as the mean ± SEM. Significance levels are indicated as follows: *P < 0.05; **P < 0.01; ***P < 0.001; ****P < 0.0001.

### Key resources table

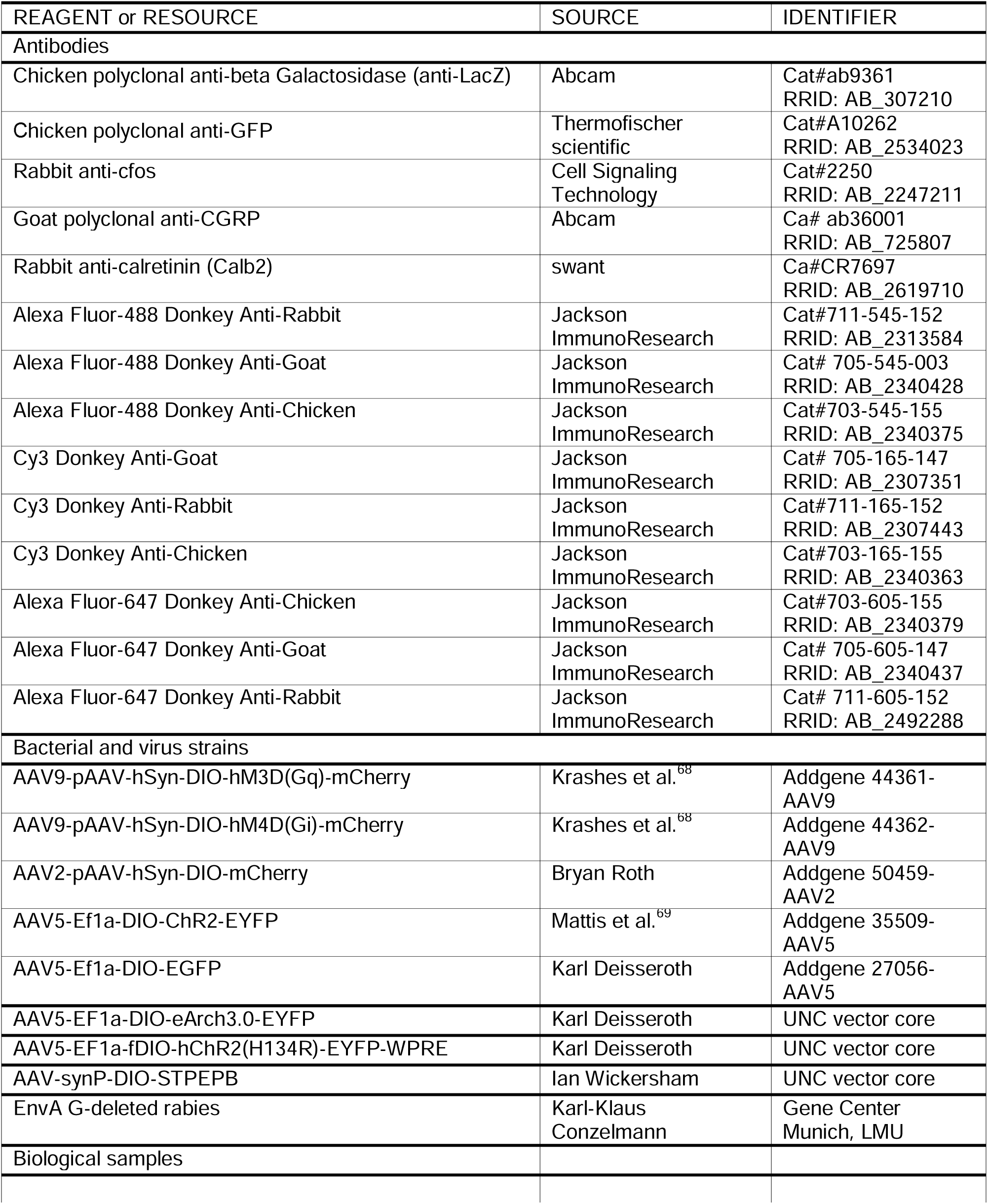
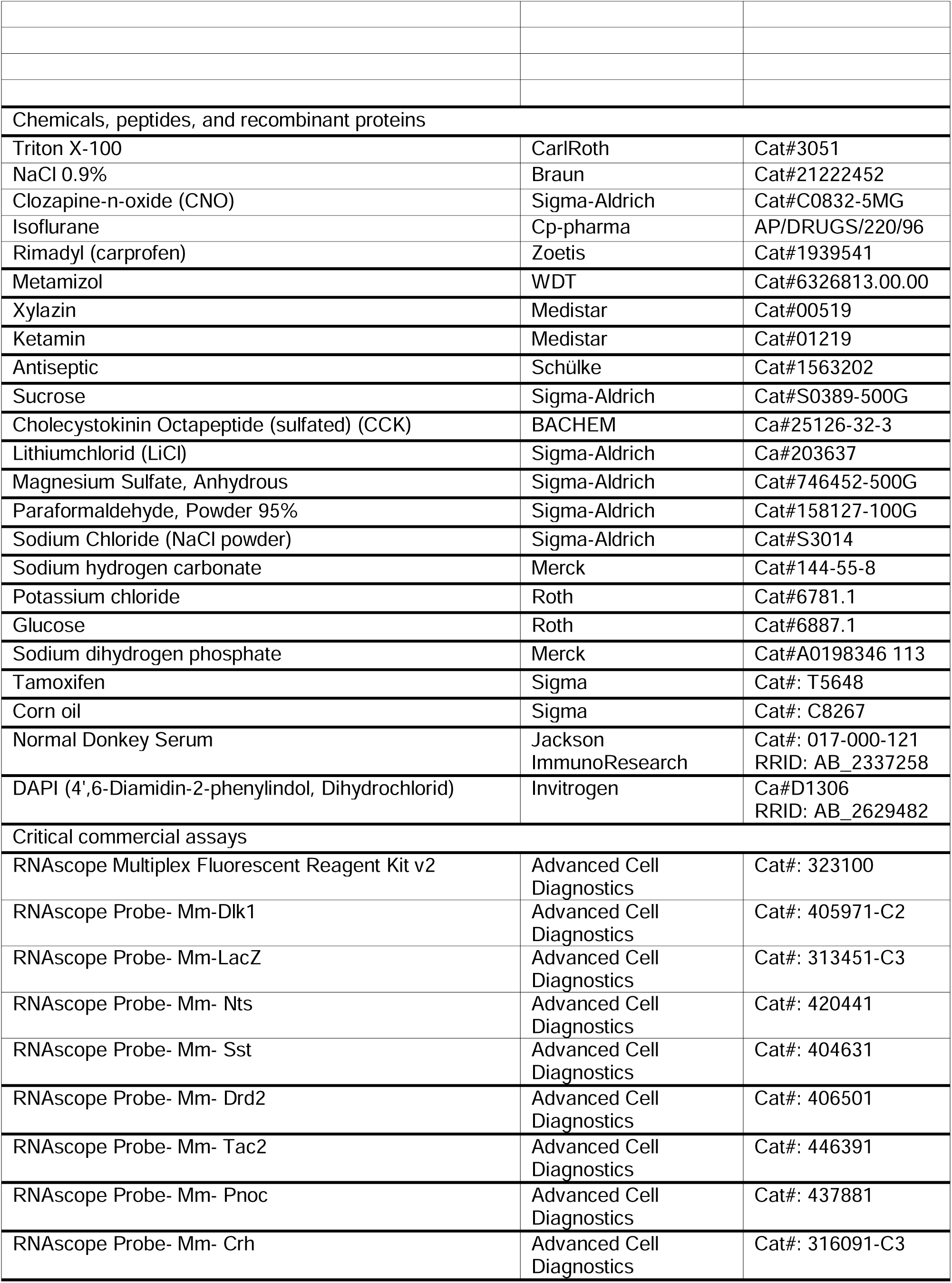
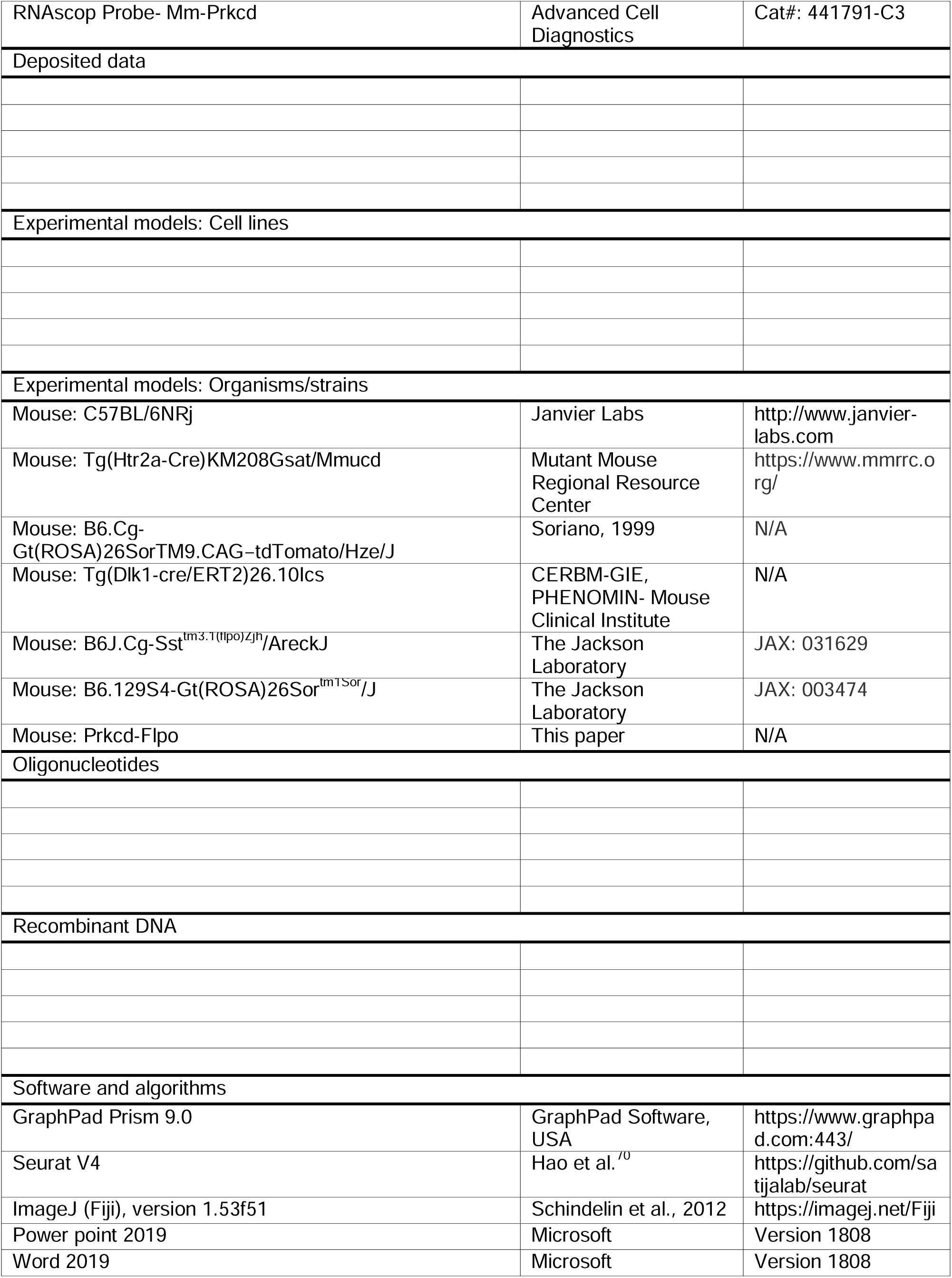
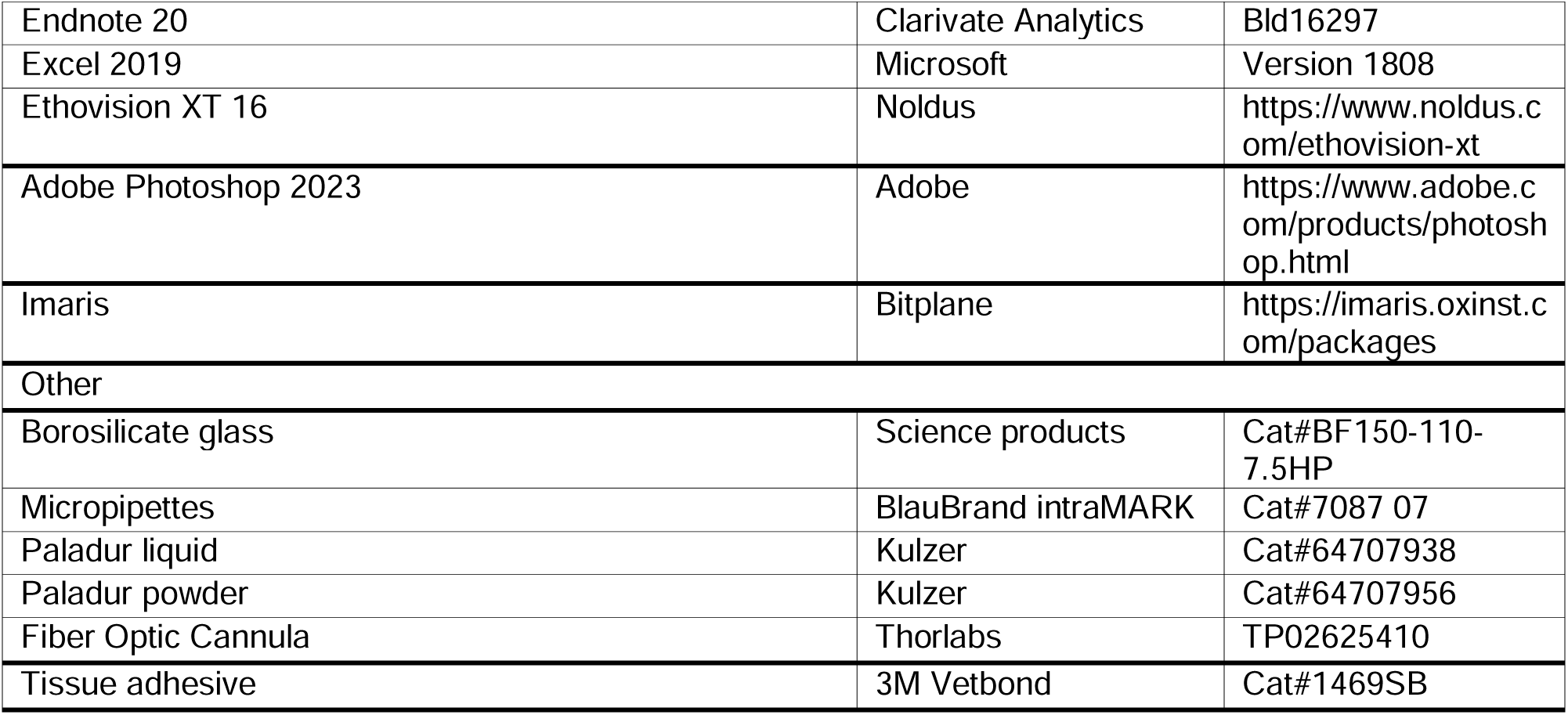

