## Supplementary Figures for "Nausea-induced suppression of feeding is mediated by central amygdala Dlk1 expressing neurons"

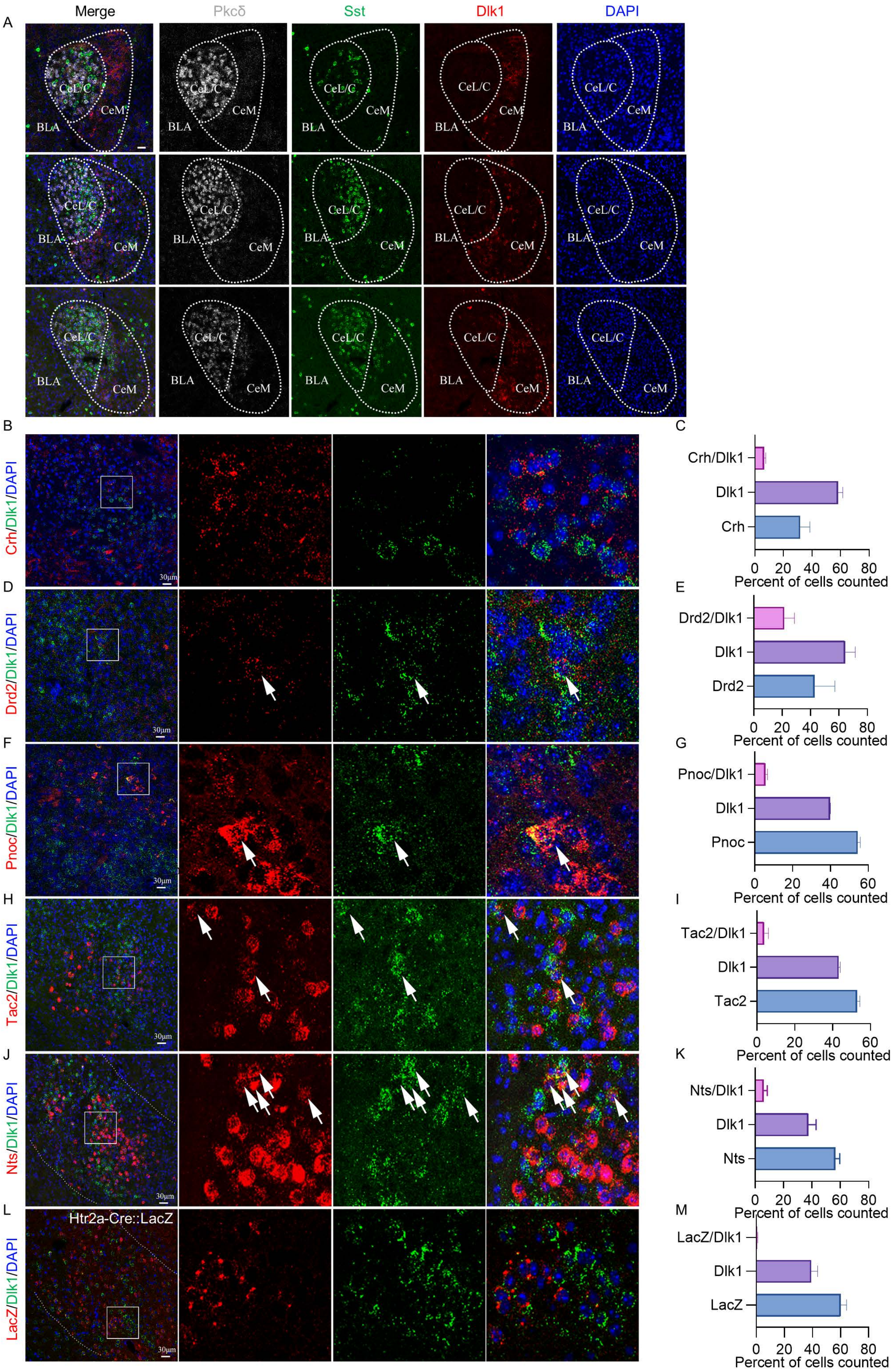


**Supplementary Figure S1**

**Supplementary Figure S1 related to Figure1:**

**Overlap between CeA$^{DLK1}$ neurons and other CeA cell types.**

**A** Representative images of *Dlk1* (red), *Sst* (green) and *Pkcδ* (grey) mRNA expression in the CeA. Scale bar, 30 µm.

**B** Representative image of *Crh* (red) and *Dlk1* (green) mRNA expression in the CeA. Scale bar, 30 µm.

**C** Quantification of Crh, Dlk1 and double-positive cell counts (3 sections per mouse, 3 mice; error bars represent SEM)

**D** Representative image of *Drd2* (red) and *Dlk1* (green) mRNA expression in the CeA. Scale bar, 30 µm.

**E** Quantification of Drd2, Dlk1 and double-positive cell counts (3 sections per mouse, 3 mice; error bars represent SEM).

**F** Representative image of *Pnoc* (red) and *Dlk1* (green) mRNA expression in the CeA. Scale bar, 30 µm.

**G** Quantification of Pnoc, Dlk1 and double-positive cell counts. (3 sections per mouse, 3 mice; error bars represent SEM).

**H** Representative image of *Tac2* (red) and *Dlk1* (green) mRNA expression in the CeA. Scale bar, 30 µm.

**I** Quantification of Tac2, Dlk1 and double-positive cell counts. (3 sections per mouse, 3 mice; error bars represent SEM).

**J** Representative image of *Nts* (red) and *Dlk1* (green) mRNA expression in the CeA. Scale bar, 30 µm.

**K** Quantification of Nts, Dlk1 and double-positive cell counts. (3 sections per mouse, 3 mice; error bars represent SEM).

**L** Representative image of Htr2a-Cre::LacZ mouse immunostained for β-galactosidase (red) and ISH for *Dlk1* (green) mRNA in the CeA. Scale bar, 30 µm.

**M** Quantification of β-galactosidase, Dlk1 and double-positive cell counts. (3 sections per mouse, 3 mice; error bars represent SEM).

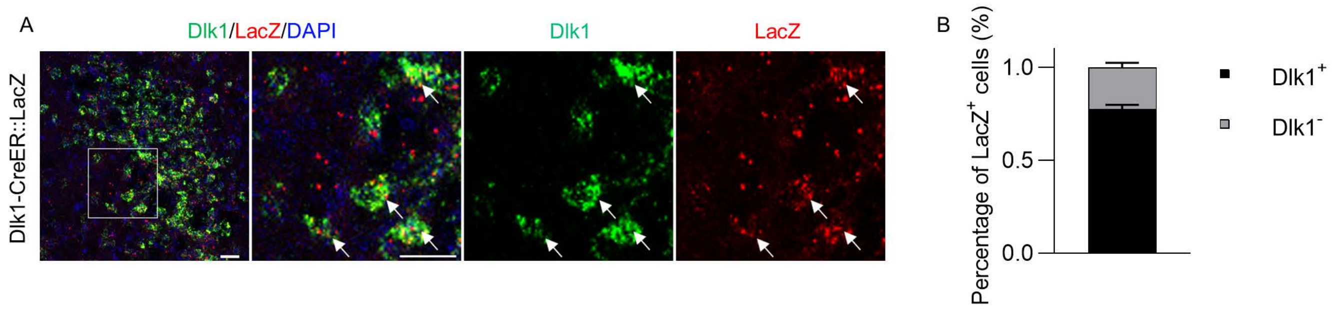


**Supplementary Figure S2 related to Figure1:**

**Dlk1-CreER mouse has a high-fidelity of Cre expression.**

**A** Representative image through the CeA of a Dlk1-CreER mouse expressing a floxed-lacZ reporter and labeled for *LacZ* and *Dlk1* mRNAs. White box indicates the location of the high-magnification panel on the right. Arrow indicate LacZ$^+$ cells that co-express Dlk1. Scale bar, 30 µm.

**B** Quantification of LacZ$^+$ neurons in the CeA express Dlk1 (n = 3 mice; error bars represent SEM).

**Supplementary Figure S2**

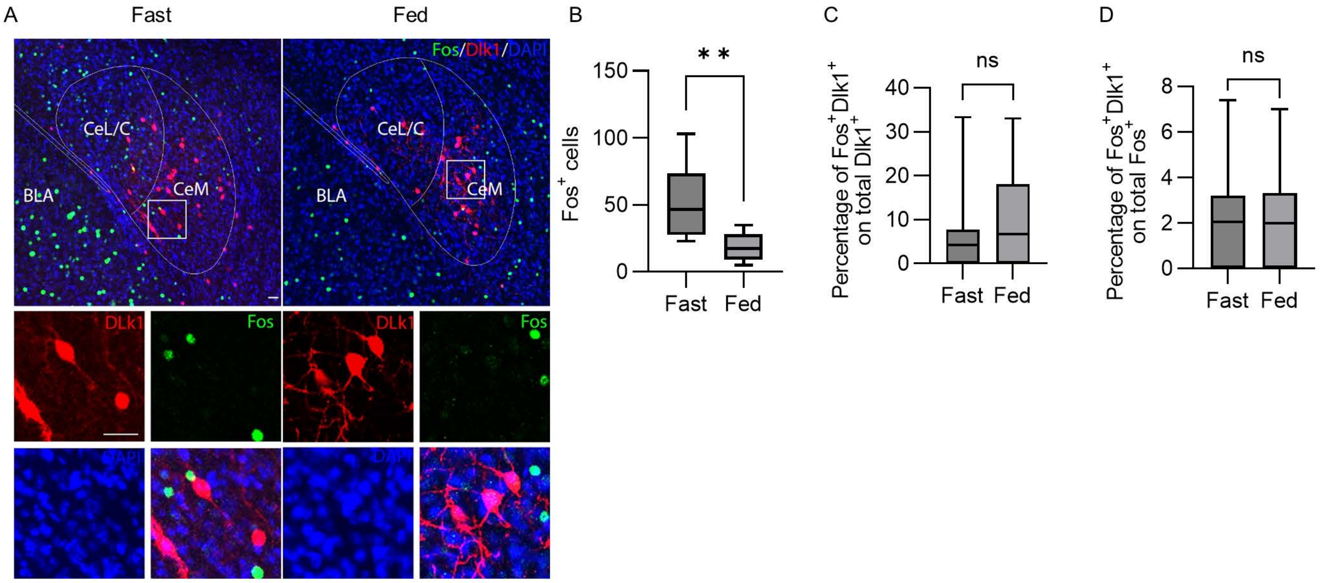


**Supplementary Figure S3 related to Figure2:**

**CeA$^{Dlk1}$ neurons are not activated by satiety.**

**A** Representative images of c-Fos immunostainings (green) in CeA of Dlk1-CreER::Ai9 tdTomato mice (red) under 20h fasted or Ad libitum fed condition. White box indicates the location of the high-magnification panel on the bottom. Scale bars, 30 µm.

**B** Percentages of c-Fos$^{+}$ cells in CeA after the indicated treatments. Data in panels B-D were analyzed by Two-tailed unpaired t test, P = 0.0033.

**C** Percentages of c-Fos$^{+}$/Dlk1$^{+}$ cells among Dlk1$^{+}$ cells in CeA after the indicated treatments. P = 0.4532.

**D** Percentages of c-Fos$^{+}$/Dlk1$^{+}$ cells among c-Fos$^{+}$ cells in CeA after the indicated treatments. P = 0.9872.

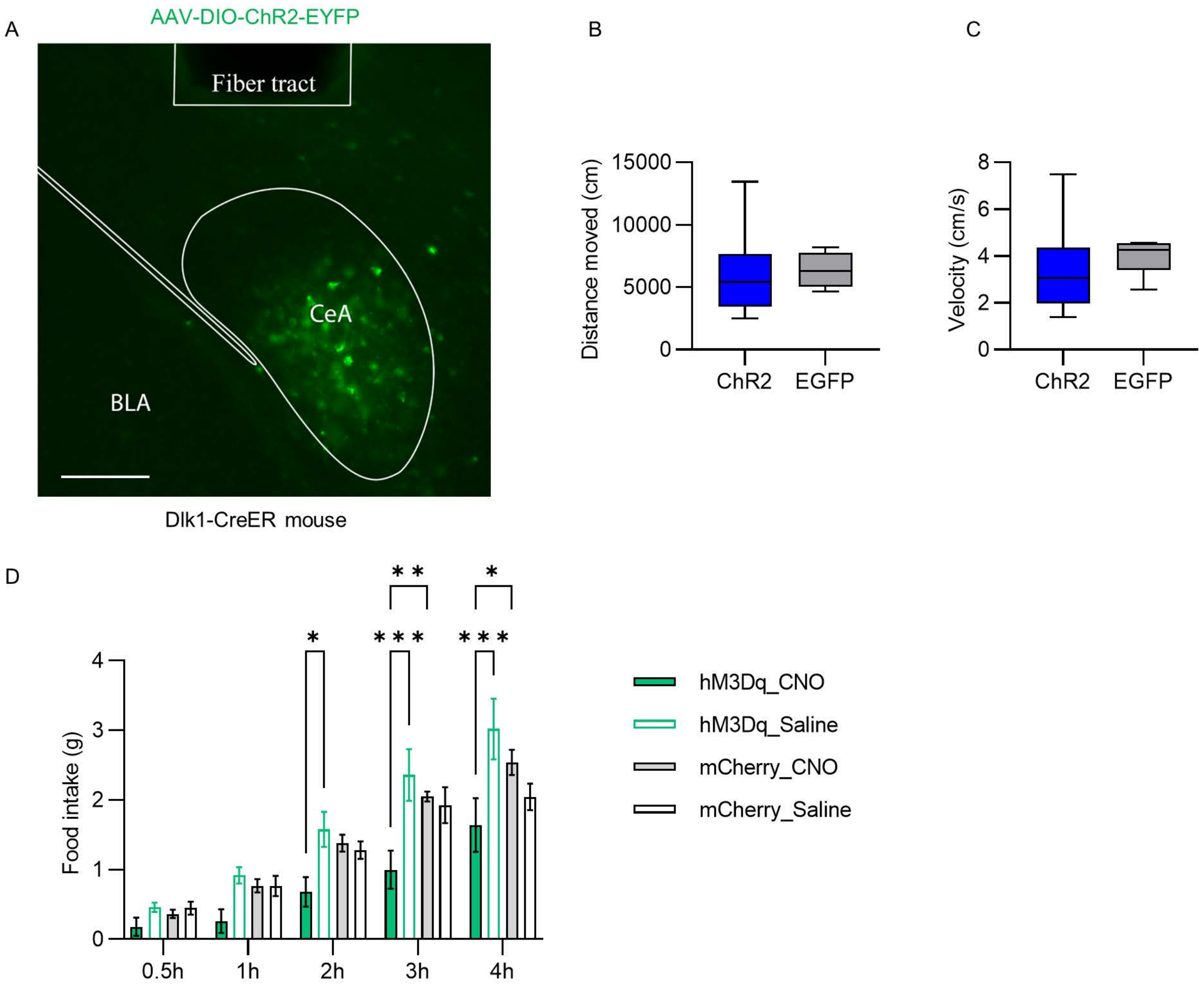


**Supplementary Figure S4 related to Figure3:**

**Activation of CeA$^{Dlk1}$ neurons inhibits feeding.**

**A** Representative histological image of ChR2 expression in the CeA with optic fiber tract. Scale bar, 100 µm

**B** Distance moved by animals expressing control protein EGFP or ChR2 during feeding test. Unpaired t test. P = 0.6242.

**C** The velocity of animals expressing control protein EGFP or ChR2 during feeding test. Unpaired t test. P = 0.2756.

**D** Cumulative food intake of satiated Dlk1-CreER animals expressing the excitatory DREADD hM3Dq (pAAV-hSyn-DIO-hM3Dq-mCherry) virus or control (pAAV-hSyn-DIO-mCherry) virus in CeA, after i.p. injections of saline and CNO (2 mg/Kg). Two-way ANOVA, *p<0.05, **p<0.01, ***p<0.001, n=5 mice per group.

**Supplementary Figure S4**

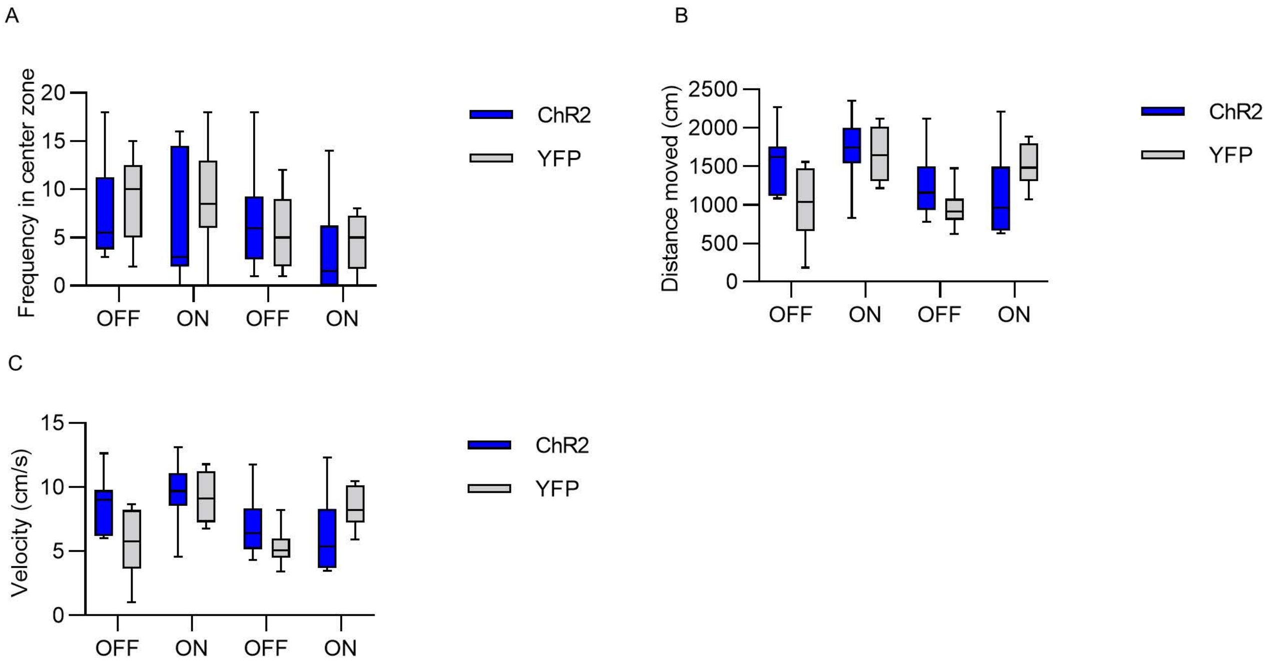


**Supplementary Figure S5 related to Figure4:**

**Activation of CeA$^{Dlk1}$ neurons does not induce anxiety.**

**A** Frequency of visits of the central zone by animals expressing control protein EGFP (n=6) or ChR2 (n=10) in the OF task. Unpaired t test. P(OFF1) = 0.5832. P(ON1) = 0.3865. P(OFF2) = 0.6618. P(ON2) = 0.5325.

**B** Distance moved by animals expressing control protein EGFP or ChR2 in the OF task. Unpaired t test. P(OFF1) = 0.1072. P(ON1) = 0.8113. P(OFF2) = 0.4011. P(ON2) = 0.4011.

**C** The velocity of animals expressing control protein EGFP or ChR2 in the OF task. Unpaired t test. P(OFF1) = 0.1088. P(ON1) = 0.8264. P(OFF2) = 0.3928. P(ON2) = 0.3928.

**Supplementary Figure S5**

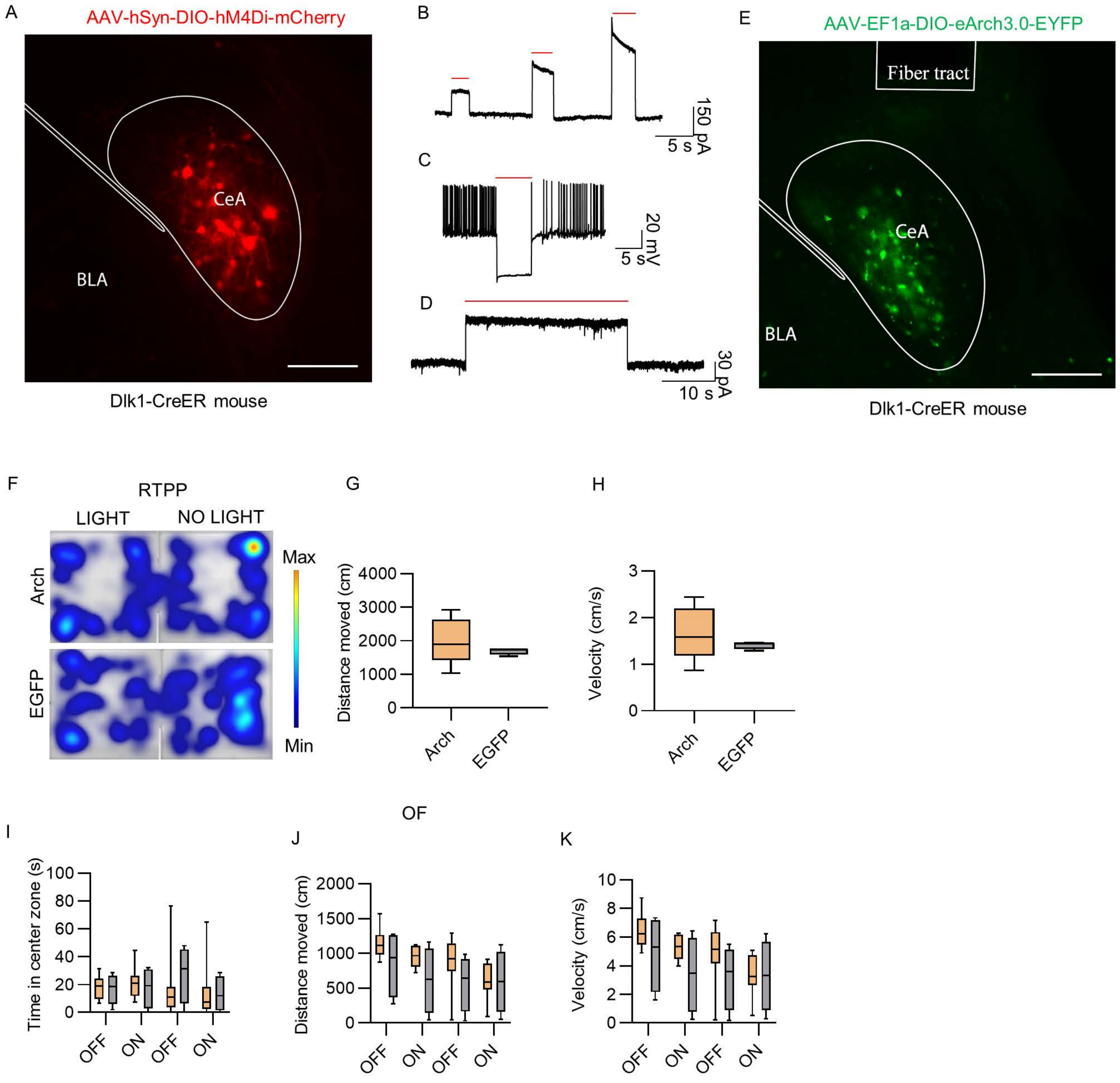


**Supplementary Figure S6**

**Supplementary Figure S6 related to Figure5:**

**Silencing $CeA^{Dlk1}$ neurons is neither rewarding nor anxiogneic.**

**A** Representative histological image of hM4Di expression in the CeA. Scale bar, 100 μm

**B** Photoinhibition in voltage clamp (-70 mV) of $CeA^{Dlk1}$ neurons using different intensities of red light (% intensity: 10, 20, 30).

**C** Photoinhibition of $CeA^{DLK1}$ neurons suppressed current injection-induced firing.

**D** Stable photoinhibition with continuous red light (30 s) of $CeA^{DLK1}$ neurons**.**

**E** Representative histological image of Arch expression in the CeA with optic fiber tract. Scale bar, 100 μm

**F** Representative images of heatmaps depicting the time spent in the RTPP task.

**G** Distance moved in the RTPP task by animals expressing control protein EGFP (n=4) or Arch (n=9). Two-tailed unpaired t test. P = 0.3721

**H** The velocity of animals expressing control protein EGFP or Archin the RTPP task. Unpaired t test. P = 0.3721.

**I** Cumulative duration in the central zone of the arena by animals expressing control protein EGFP (n=4) or Arch (n=9) in the OF task.. Unpaired t test. P(OFF1) = 0.9819. P(ON1) = 0.9716. P(OFF2) = 0.8987. P(ON2) = 0.9819.

**J** Distance moved in the OF task by animals expressing control protein EGFP or Arch.Unpaired t test. P(OFF1) = 0.1443. P(ON1) = 0.0677. P(OFF2) = 0.2177. P(ON2) = 0.9085.

**K** The velocity of animals expressing control protein EGFP or Arch in the OF task. Unpaired t test. P(OFF1) = 0.1622. P(ON1) = 0.0677. P(OFF2) = 0.2177. P(ON2) = 0.9085.

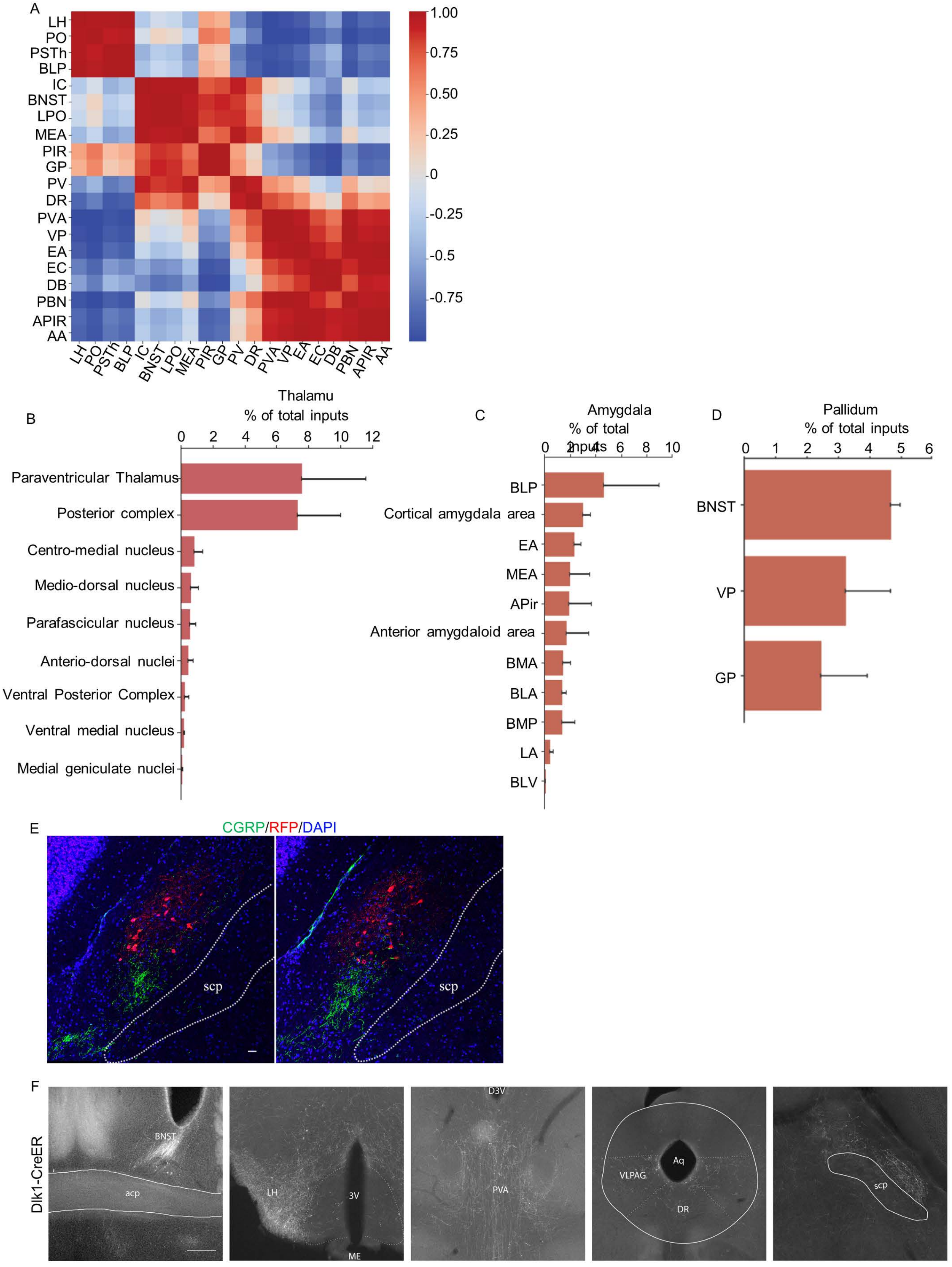


**Supplementary Figure S7**

**Supplementary Figure S7 related to Figure6:**

**Monosynaptic afferent and efferent connections of CeA$^{Dlk1}$ neurons.**

**A** Clustering on Pearson's pairwise correlation coefficients between the most input regions counted from Dlk1-CreER-tracing experiments. The color scale indicates the degree of correlation (n=3 mice).

**B** Brain regions in the thalamus projecting to CeA$^{Dlk1}$ neurons (n = 3 mice; error bars represent SEM). Posterior complex includes: PO, posterior thalamic nuclear group, triangular Part (PoT), posterior intralaminar thalamic nucleus (PIL).

**C** Brain regions in the amygdala projecting to CeA$^{Dlk1}$ neurons (n = 3 mice; error bars represent SEM).

**D** Brain regions in the pallidum projecting to CeA$^{Dlk1}$ neurons (n = 3 mice; error bars represent SEM).

**E** Representative images of PBN cells that project to CeA$^{Dlk1}$ neurons (red) and CGRP expression across two different positions. Scale bar, 30 µm.

**F** Representative images of brain areas innervated by CeA$^{Dlk1}$ neurons. Scale bar, 250 µm

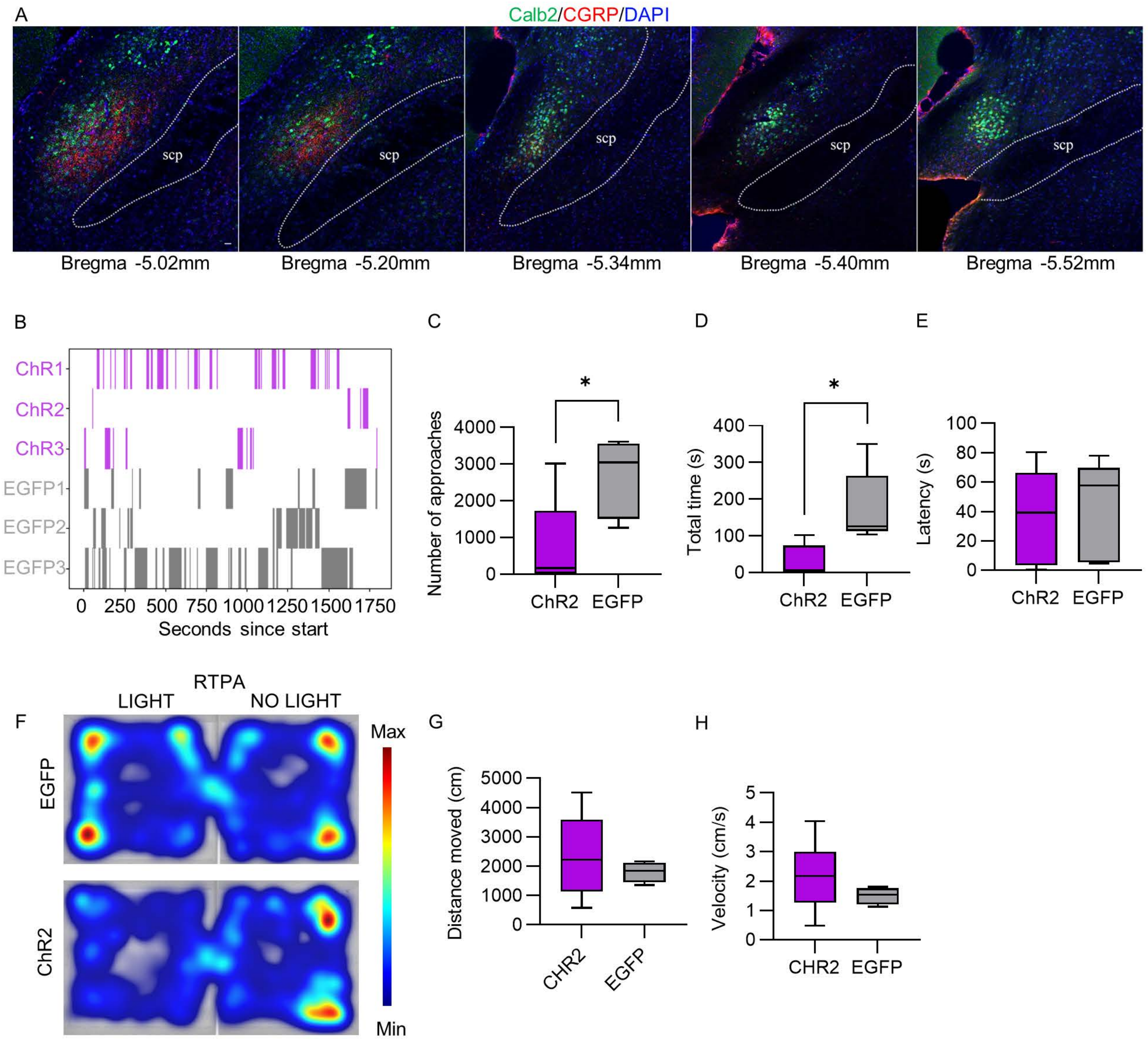


**Supplementary Figure S8**

**Supplementary Figure S8 related to Figure7:**

**The anorexigenic activity of $CeA^{Dlk1}$ neurons is mediated by projections to the PBN.**

**A** Representative images of CGRP (red) and Calb2(green) expression in the PBN across different positions, Scale bar, 30 µm.

**B** Raster plot of food approaches of 20h fasted individual mice from the photoactivated ChR2 and EGFP groups.

**C** Numbers of food approaches of 20h fasted photoactivated ChR2 and EGFP groups. Two-tailed unpaired t test, P = 0.0338.

**D** Total time of food approach of 20h fasted photoactivated ChR2 and EGFP groups. Two-tailed unpaired t test, P = 0.0188.

**E** Latency of the first approach to food of 20h fasted photoactivated ChR2 and EGFP groups. Two-tailed unpaired t test, P = 0.5048.

**F** Representative images of heatmap depicting the time spent in the RTPP task.

**G** Distance moved in the RTPA task by animals expressing control protein EGFP or ChR2. Two-tailed unpaired t test. P = 0.3783.

**H** The velocity of animals expressing control protein EGFP or ChR2 in the RTPA task. Unpaired t test. P = 0.2076.
